# Defective Function of α-Ketoglutarate Dehydrogenase Exacerbates Mitochondrial ATP Deficits during Complex I Deficiency

**DOI:** 10.1101/2020.01.09.900514

**Authors:** Gerardo G. Piroli, Allison M. Manuel, Richard S. McCain, Holland H. Smith, Oliver Ozohanics, Sara Mellid, William E. Cotham, Michael D. Walla, Alberto Cascón, Attila Ambrus, Norma Frizzell

## Abstract

The *NDUFS4* knockout (KO) mouse phenotype resembles the human Complex I deficiency Leigh Syndrome. The irreversible succination of protein thiols by fumarate is increased in select regions of the *NDUFS4* KO brain affected by neurodegeneration, suggesting a mechanistic role in neurodegenerative decline. We report that dihydrolipoyllysine-residue succinyltransferase (DLST), a component of the α-ketoglutarate dehydrogenase complex (KGDHC) of the tricarboxylic acid (TCA) cycle, is succinated in the *NDUFS4* KO brain. Succination of DLST reduced KGDHC activity in the brainstem (BS) and olfactory bulb (OB) of KO mice. The defective production of KGDHC derived succinyl-CoA resulted in decreased mitochondrial substrate level phosphorylation (SLP), further aggravating the OXPHOS ATP deficit. Protein succinylation, an acylation modification that requires succinyl-CoA, was reduced in the KO mice. Modeling succination of a cysteine in the spatial vicinity of the DLST active site or introduction of succinomimetic mutations recapitulates these metabolic deficits. Our data demonstrate that the biochemical deficit extends beyond impaired Complex I assembly and OXPHOS deficiency, functionally impairing select components of the TCA cycle to drive metabolic perturbations in affected neurons.

## Introduction

Leigh syndrome is a mitochondrial disease predominantly caused by single gene defects in the oxidative phosphorylation (OXPHOS) machinery or the pyruvate dehydrogenase complex. It is characterized by bilateral necrotizing lesions of the basal ganglia and brainstem (BS), lactic acidosis, ataxia, seizures, and respiratory failure (Lake et al., 2016). Leigh syndrome is associated with mutations in at least 23 genes that lead to mitochondrial Complex I deficiency (Lake et al., 2016), with mutations in *NDUFS4*, a gene that encodes the small assembly protein NADH dehydrogenase (ubiquinone) iron-sulfur protein 4 (Ndufs4) (Calvaruso et al., 2011; Ortigoza-Escobar et al., 2016), among the most frequent.

The homozygous *NDUFS4* knockout mouse (Ndufs4 KO) recapitulates many biochemical and clinical aspects of Leigh syndrome, including lactic acidosis, BS degeneration, motor retardation and fatal respiratory failure at 8 weeks after birth; the neuropathology also extends to the olfactory bulb (OB) and some cerebellar nuclei (Kruse et al., 2008; Quintana et al., 2010). Selective deletion of Ndufs4 in glutamatergic neurons leads to BS inflammation, motor and respiratory deficits and early death, whereas ablation of Ndufs4 in GABAergic neurons causes basal ganglia inflammation, hypothermia, and severe epileptic seizures. This highlights the combined contributions of different neuronal populations to the complex pathology in this model (Bolea et al., 2019).

While the genetic defects underlying Leigh syndrome are well documented, the biochemical mechanisms linking the bioenergetic deficit to the loss of select neuronal populations remains ill defined. Increased oxidative stress has been implicated in the pathogenesis of the Ndufs4 KO (Quintana et al. 2010; de Haas et al., 2017, Kelly et al., 2023), attributed in part to elevated local hyperoxia and increased free iron (Jain et al., 2019, Kelly et al. 2023). Reductive stress is a well-established component of impaired OXPHOS during mitochondrial disease (Titov et al., 2016, Sharma et al., 2021). The ratio of NADH/NAD^+^ is elevated in the Ndufs4 KO model (Lee et al. 2019), and we have described that loss of Ndufs4 contributes to alterations in the NAD^+^-dependent tricarboxylic acid (TCA) cycle. Specifically, we have detected elevated levels of fumarate-derived protein succination in the pathologically lesioned regions of the Ndufs4 KO brain, namely brainstem and olfactory bulb (Piroli et al. 2016). Protein succination is a covalent chemical modification of protein cysteine thiols following non-enzymatic modification by fumarate, generating S-2-succinocysteine (2SC) (Figure S1A, Alderson et al., 2006; Frizzell et al., 2009; Merkley et al., 2014), a modification that contributes to reduced protein function (Manuel et al., 2017; Piroli et al. 2014; Ternette et al. 2014). Thiol reactivity with protonated fumarate is enhanced in an acidic microenvironment (Kulkarni et al. 2019), and the Ndufs4 KO is a well-characterized model of metabolic acidosis. While abundant succination of tubulin is endogenously present in all neurons due to the long-lived axonal microtubules, we confirmed increased succination of select proteins in lesioned Ndufs4 KO brain regions such as the vestibular nucleus, e.g., mitochondrial voltage-dependent anion channels (VDAC) 1 and 2 (Piroli et al., 2016).

In the current study, we demonstrate that mitochondrial dihydrolipoyllysine-residue succinyltransferase (DLST), a component of the α-ketoglutarate dehydrogenase complex (KGDHC), is succinated in the Ndufs4 KO brain. KGDHC is a significant rate limiting step of the TCA cycle in the brain (Sheu and Blass, 1999), suggesting that impaired activity of this complex would have pronounced effects when OXPHOS is defective, indeed homozygous loss of DLST is embryonically lethal (Kiss et al., 2013). In humans, mutation of *OGDH* in two siblings resulted in a ∼50% decrease in OGDH activity and manifested as a neurological mitochondrial disease (Yap et al., 2020). Two recent studies profiling regional brain differences by global metabolomic profiling reinforce the central importance of the KGDHC axis in the Ndufs4 KO mouse model. Johnson et al. have documented defective α-ketoglutarate metabolism to glutamate and glutamine. Moreover, they demonstrate that the beneficial effect of rapamycin treatment in this model was via the augmentation of α-ketoglutarate (Johnson et al. 2020). Terburgh et al. also describe in detail defective α-ketoglutarate-derived succinate and glutamate production and increased branched chain amino acids in parallel with decreased BCAA-CoAs, which would yield even less entry of anaplerotic succinyl CoA to the defective TCA cycle (Terburgh et al. 2021a). Here, we report the striking functional impact of DLST succination and defective KGDHC activity on distinct mitochondrial metabolic processes, providing a mechanistic link between the documented metabolomic defects in this model and the region-specific brain pathology. These results biochemically define how mitochondrial ATP deficiency is exacerbated beyond the existing genetic Complex I bioenergetic defect.

## Results

### Dihydrolipoyllysine-residue succinyltransferase is succinated in the Ndufs4 KO mouse brain

Protein succination is selectively increased in the brainstem of the Ndufs4 KO mouse versus WT (Figure 1A). We have previously described the abundant and consistent succination of tubulin isoforms in the brainstem (BS) and the olfactory bulb (OB) of both WT and Ndufs4 KO mice (Piroli et al., 2016). Using immunoblotting with an anti-2SC antibody the pronounced succinated tubulin band is resolved at ∼50 kDa in BS protein homogenates, appearing slightly larger and expanding to a lower MW in the KO mice vs. WT (Figure 1A). To determine if additional proteins were succinated at this molecular weight and masked by the intensity of tubulin succination, we performed a tubulin polymerization assay that we had previously employed (Piroli et al., 2014) in which the initial high speed centrifugation pellet is substantially depleted of tubulin. Immunoblotting with anti-2SC following this tubulin depletion approach allowed the resolution of any residual succinated tubulin from a new succinated protein that was only present in the KO mouse brainstem (Figure 1B, arrow in 2SC panel). A longer exposure shows enrichment of this ∼48-50 kDa succinated band following depletion of cytosolic tubulin by polymerization (Figure S1B, arrow in 2SC panel). In a separate experiment to determine where succinated proteins might be localized, we isolated gliosomes and synaptosomes from WT and Ndufs4 KO mouse BS to determine where succinated proteins might be localized. Glial fibrillary acidic protein (GFAP), an astrocytic marker, was enriched in the gliosomes, whereas the presence of the neuronal presynaptic marker synaptophysin was prominent in the synaptosomes (Figure 1C, GFAP and synaptophysin panels). A final pellet from these enrichments contains other organelles including the mitochondria, and we noted the outer mitochondrial membrane marker VDAC2 was present in the original tissue homogenate and the synaptosomes but was most abundant in this remaining pellet (Figure 1C, VDAC2 panel). Overall, the pellets were rich in mitochondria but devoid of cytosolic, synaptic and astroglial markers. After probing with anti-2SC antibody, the succinated band at ∼48-50 kDa was strikingly enriched in the pellet fraction of Ndufs4 KO but not in WT mice (Figure 1C, arrow in 2SC panel). Cytosolic tubulin depletion facilitated the detection of the succinated band at ∼48-50 kDa (Figure 1C, α-tubulin panel), as the abundant presence of succinated tubulin both in WT and KO tissue homogenates, gliosomes and synaptosomes overwhelms the detection of this ∼48-50 kDa band (Figure 1C, 2SC panel). Similar results were obtained when we performed the same enrichment with the cerebellum (CB), another affected brain region in Ndufs4 KO mice (Figure S1C, D).

**Figure 1:**
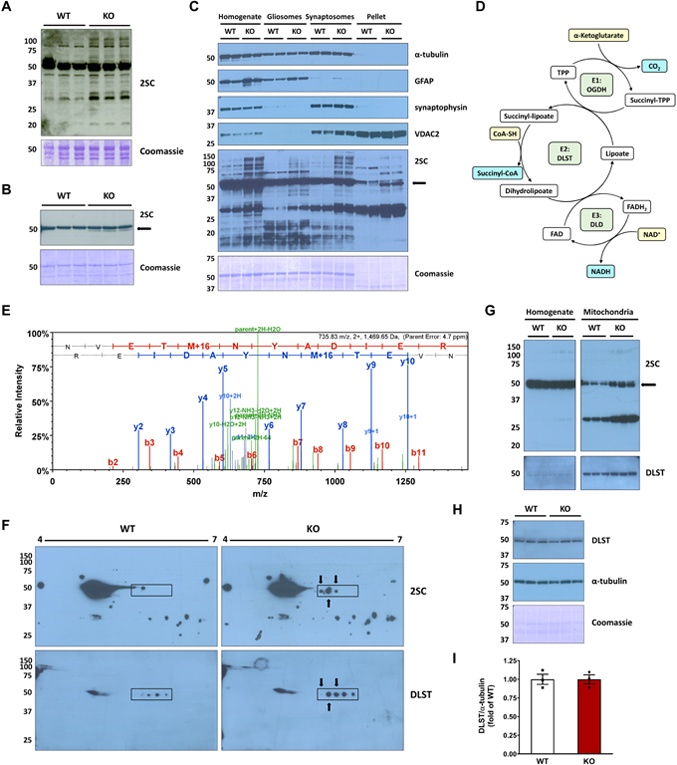
Identification of dihydrolipoyllysine-residue succinyltransferase (DLST) succination in the brainstem (BS) of the Ndufs4 KO mouse. *A*, Total levels of protein succination (2SC) in brainstem (BS) homogenates from WT and Ndufs4 KO mice, note the abundance of succinated tubulin in WT and KO at ∼50kDa. Ndufs4 KO BS show enrichment of several additional succinated proteins vs. WT mice. *B*, Detection of a distinct succinated band at ∼48-50 kDa (arrow) in Ndufs4 KO BS following cytosolic tubulin depletion. *C*, Characterization of the purity of total tissue protein homogenates, gliosomes, synaptosomes, and a mitochondria-rich pellet fraction obtained from BS by analysis of α-tubulin, GFAP, synaptophysin and VDAC2 expression. A ∼48-50 kDa succinated band is present in the pellet fractions (2SC panel, KO lanes, arrow). *D*, Structure and function of the α-ketoglutarate dehydrogenase complex (KGDHC). *E,* MS/MS spectrum of the peptide NVETM^ox^NYADIER corresponding to DLST in a ∼48-50 kDa band isolated from the mitochondria-rich pellets used in *C*. *F*, Co-localization (arrows) of succinated spots with DLST-immunoreactive spots in the region of ∼48-50 kDa and pH 5.5-6.3 (rectangular area) in Ndufs4 KO BS. *G,* Analysis of BS tissue homogenates, where tubulin precludes DLST detection, alongside purified mitochondrial fractions showing a succinated band at ∼48-50 kDa in KO lanes (2SC panel, arrow), which overlapped with DLST detection. *H*, Total DLST and α-tubulin content in BS homogenates of WT and Ndufs4 KO mice. *I*, Quantification of the DLST band intensity in *H*. See also Figure S1. In *A-C*, and *F-H*, molecular weight markers are shown on the left side. Coomassie staining was used to verify even load of the lanes. In *D*, substrates highlighted in yellow, products in light blue and the components of the complex in light green. In *I*, results expressed as mean ± SEM were compared by unpaired t test (NS).

The mitochondria-rich pellet fractions were further separated by SDS-PAGE, with parallel lanes used for anti-2SC immunoblotting or Coomassie staining, prior to band excision ∼48-50 kDa region and LC-MS/MS mass spectrometry. Several abundant mitochondrial proteins with significant XCorr scores were identified from the in-gel trypsin digests, but a specific succination site could not be assigned as the detection of endogenous levels of negatively charged succinated peptides is difficult in positive ion mode mass spectrometry. For each of these proteins we manually calculated the isoelectric point (pI) of the mature mitochondrial form (minus the signal sequence) to determine if any aligned with a distinct series of succinated spots that we repeatedly resolved following 2D-electrophoresis and immunoblotting (Figure 1F, boxed region in upper 2SC panels). Several succinated spots were present in the KO this area, whereas in the WT only one spot was present at a slightly higher molecular weight (Figure 1F, 2SC panels, WT vs. KO boxed regions). We have previously noted that this succinated spot series appeared around a pI ∼6 when resolved away from tubulin isoforms whose pI is 4.7-5.4 (Figure 1F, 2SC panels, Piroli et al., 2014, Piroli et al. 2016). The protein dihydrolipoyllysine-residue succinyltransferase (DLST), a component of the α-ketoglutarate dehydrogenase complex (KGDHC), had a computed isoelectric point (pI) ∼5.98 for the mature mitochondrial form (removing transit peptide 1-68). The KGDHC comprises several copies of three subunits (Figure 1D): 2-oxoglutarate dehydrogenase (OGDH, E1), DLST (E2), and dihydrolipoamide dehydrogenase (DLD, E3) (Reed and Oliver, 1982). A representative DLST-specific peptide NVETM^OX^NYADIER ([M+2H]^2+^: 735.8322, XCorr (+2) = 3.94) is shown in Figure 1E, other DLST peptides confirmed by mass spectrometry are summarized in Figure S1E. Applying a DLST-specific antibody to the 2-D immunoblots we detected precise co-localization with three succinated spots in the KO (Figure 1F, overlap marked by arrows in KO 2SC and DLST panels). In contrast, the four spots detected by anti-DLST do not overlap with any succinated spot in the WT (Figure 1F, WT DLST and 2SC panel). We further analyzed both BS tissue homogenates and purified mitochondrial fractions from WT and Ndufs4 KO BS to examine if the succinated protein paralleled DLST localization. Figure 1G shows a short exposure of BS tissue homogenates from KO and WT mice without detectable changes in succination (2SC panel, homogenate lanes) due to the abundance of the succinated tubulin band (Piroli et al., 2014; Piroli et al., 2016). In contrast, mitochondrial purification allowed for better resolution of the distinctly succinated ∼48-50 kDa band in the KO BS (2SC panel, mitochondria lanes, black arrow). The additional succinated band at ∼30-32 kDa had been previously identified as mitochondrial VDAC1 and 2 (Piroli et al., 2016). After stripping, a specific anti-DLST antibody showed similar levels of this protein in both WT and Ndufs4 KO mitochondria, overlapping with the succinated bands (DLST and 2SC panels, mitochondria lanes). We confirmed that total DLST protein levels do not change in the BS between genotypes, as shown in Figure 1H and 1I. In addition, we did not detect any changes in OGDH levels (E1 subunit) or other TCA cycle enzymes, indicating TCA cycle enzyme levels are not reduced in the KO model (Figure S1F). In summary, DLST is succinated by fumarate in the BS of Ndufs4 KO mice compared to WT mice.

### Succination reduces the activity of the α-ketoglutarate dehydrogenase complex, affecting substrate level phosphorylation

We hypothesized that DLST succination driven by increased fumarate might affect the functionality of the KGDHC, since components of this complex are susceptible to oxidative modification (Ambrus et al., 2009; Chinopoulos et al., 1999; Humphries and Szweda, 1998; Starkov et al., 2004). We confirmed increased fumarate concentration in the olfactory bulbs (OB) of the Ndufs4 KO mouse (96.7% greater than in WT OB, p<0.001, Figure 2A), which accumulated in parallel with increased lactate (Figure 2B). Pyruvate and malate levels were also significantly increased in the Ndufs4 KO OB (Figure S2A and D). We then measured KGDHC activity in both the OB and BS of WT and Ndufs4 KO mice and found that in both brain regions it was significantly lower in the KO mice (31.2% and 24.2% decrease in OB and BS, p<0.01 and p<0.05 respectively, Figures 2C and D). These reductions were specific for KGDHC activity in OB and BS of the Ndufs4 KO mouse, as the activity of citrate synthase, a TCA cycle enzyme commonly used to normalize the activity of other mitochondrial enzymes, was unaffected in the same brain regions (Figure S2E and F for OB and BS, respectively). A recent report has shown increases, and not a decrease, in citrate synthase activity in the affected regions of the Ndufs4 KO brain (Terburgh et al. 2021a), further demonstrating that the KGDHC is selectively impacted. Since the Ndufs4 KO mice have a metabolic acidosis, it could be expected that this would favor the conversion of any accumulated α-ketoglutarate towards L-2-hydroxyglutarate (Nadtochiy et al. 2016). Quantification of L-2-hydroxyglutarate showed a trend (P=0.055) to decrease in the OB of the Ndufs4 KO mouse (Figure 2E), whereas α-ketoglutarate concentration was unchanged (Figure 2F). Decreased 2-hydroxyglutarate has recently been documented in the Ndufs4 KO mouse (Terburgh et al. 2021b) and suggests that any α-ketoglutarate conversion to this metabolite may rapidly be redirected to supply the mitochondrial Q-pool via the activity of L-2 hydroxyglutarate dehydrogenase.

**Figure 2:**
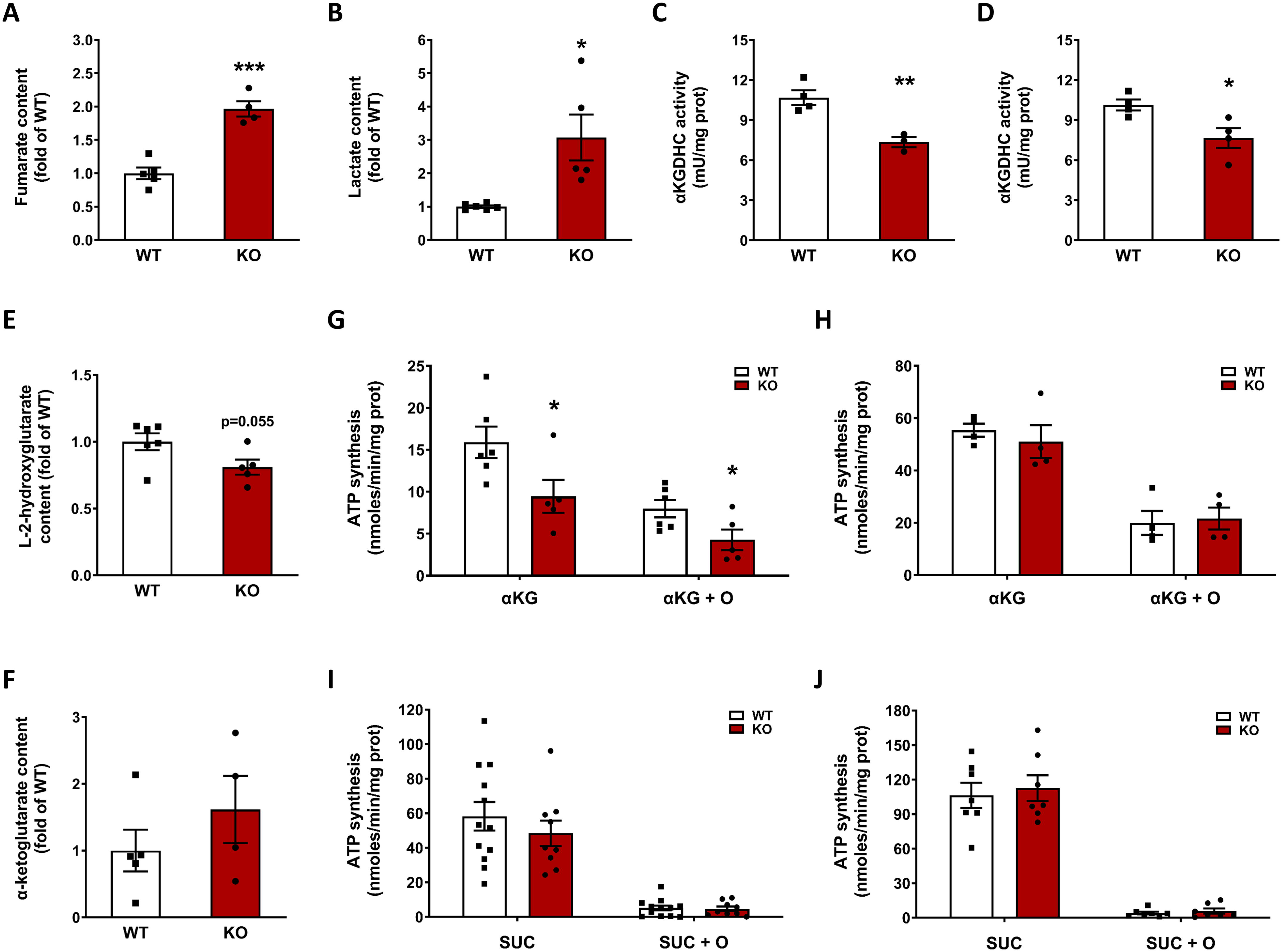
α-ketoglutarate dehydrogenase complex (KGDHC) activity is reduced as a consequence of fumarate derived protein succination. *A-B*, Fumarate (*A*) and lactate (*B*) contents in the olfactory bulb (OB) of Ndufs4 KO and WT mice. *C-D*, KGDHC activity in the OB (*C*) and BS (*D*) of Ndufs4 KO and WT mice. *E-F,* L-2-hydroxyglutarate (*E*) and α-ketoglutarate (*F*) contents in the OB of Ndufs4 KO and WT mice. *G-H*, Mitochondrial total ATP synthesis (no oligomycin) and substrate level phosphorylation (SLP, +8 µM oligomycin=O) in the OB (*G*) and the brainstem (BS) (*H*) in the presence of 5 mM α-ketoglutarate (αKG) as respiratory substrate. *I-J*, Same as in *G-H* but using 5 mM succinate (SUC) as respiratory substrate.

Since decreased KGDHC activity would lead to reduced formation of succinyl CoA, this in turn would likely decrease the conversion of succinyl-CoA into succinate in the TCA cycle. This step is catalyzed by the enzyme succinyl-CoA ligase (SUCLA), which is responsible for the synthesis of GTP or ATP by substrate level phosphorylation (SLP). In rodents and human brain, succinyl-CoA ligase preferentially produces ATP, whereas in anabolic tissues the main product is GTP (Lambeth et al., 2004; Ostergaard, 2008). We measured total and SLP-linked ATP synthesis in mitochondria isolated from both OB and BS of WT and Ndufs4 KO mice. Under the conditions used, total ATP synthesis represents the sum of ATP production by OXPHOS and SLP, and the residual ATP synthesis in the presence of the ATP synthase inhibitor oligomycin represents SLP (Komlódi and Tretter, 2017). Using α-ketoglutarate as a substrate, total ATP synthesis was decreased by 42.5% (p<0.05), and SLP was decreased by 48.3% (p<0.05) in the OB of the KO mice (Figure 2G). These data demonstrate that, in addition to the Complex I defect in OXPHOS derived ATP production, the generation of TCA cycle derived intramitochondrial ATP is also compromised. In contrast, we observed no significant differences in total ATP synthesis and SLP in the BS of Ndufs4 KO vs. WT mice (Figure 2H). This may reflect the heterogeneity of the total mitochondrial protein used fraction, in the BS select nuclei are more affected by pathology, e.g., the vestibular nucleus. When succinate was used as a respiratory substrate, no differences in total ATP synthesis were observed for OB (Figure 2I) and BS (Figure 2J). As expected, succinate did not support SLP irrespective of the genotype and brain region (Figure 2I and J) (Komlódi and Tretter, 2017). Analysis of the succination profile of the isolated mitochondria used for these SLP analyses showed a succinated band at ∼48-50 kDa that was only present Ndufs4 KO mice (Figure S2H), underscoring the selectivity of succination.

To better understand the impact of DLST succination on the KGDHC functionality, we performed molecular dynamics simulation of human DLST (hDLST) based on the recently characterized 3D structure (Nagy et al., 2021), and preliminary 3D assessment of this recent model demonstrated that Cys178 was located in the spatial vicinity of the active site. The mature forms of both murine and human DLST contain 2 cysteines and we assumed that the striking homology of human and mouse DLST would allow us to draw conclusions applicable to our mouse model (Figure S3A). We focused on the effects of Cys178 succination (2SCys178) by modeling the covalent succination modification at this site. The possible role of a second cysteine succination (i.e., Cys37) on the functionality of hDLST could not be evaluated due to the very high flexibility of the region before Asn161 that leads to unstable simulation. Molecular dynamics simulation of hDLST showed that the interaction pattern was altered in select regions of the protein when Cys178 was succinated. For instance, the Arg358-Asp356 salt bridge was perturbed due to the immediate spatial vicinity of the succinated Cys178 residue, leading to a new ionic interaction of this residue with Arg358 (distance of 2.71 Å, Figure S3C). In addition, in the unmodified hDLST, Ala179 and Asp356 (distance of 2.92 Å, Figure S3C) formed a H-bond that stabilized the relative positions of His357 (and the loop to which His357 belongs), the β_B_ and β_J_ strands; this is important for proper channeling of the Coenzyme A (CoA) and the succinyl-dihydrolipoamide moiety (SLAM) substrates (Nagy et al., 2021, see structure in Figure 3A and 3B). Conversely, succination of Cys178 resulted in the disruption of this H-bond, as the average distance between Ala179 and Asp356 increased to 4.81 Å (Figure S3C); this change affects the three chains in the hDLST trimer in a similar manner (Figure S3D). Rearrangement of the local interaction pattern modified the positions of the aforementioned beta strands and increased the distance between channel-forming side chains, which might lead to less efficient coordination of the CoA and SLAM substrates, resulting potentially in increased K_M_ values. The position/orientation of His357, part of the DLST active site, was also changed in the succinated hDLST (Figure 3A) and may alter this residue’s capacity to either deprotonate the CoA substrate or stabilize the tetrahedral oxyanionic catalytic intermediate (Nagy et al., 2021). Unfortunately, we could not draw reliable conclusions regarding the interactions between the monomers, since the dynamic simulations were found to be too unstable; this might be a result of omitting residues 150-160, which participate in the stabilization of the trimeric structure. Overall, the observed structural changes were mostly consistent for the three protein chains, showing the reliability of the simulation. The hDLST regions most affected by succination encompass the first 12 and last 8 residues studied, which in the unmodified structure are stable helical structures (Figure S3E). Although the overall active site region showed shifts lower than 0.5 Å on average in the succinated structure, select residues involved in substrate coordination showed much greater shifts (Figure 3B). This simulation work strongly supports that succination of Cys178 impacts both catalytically important active site residues and the substrate-binding channels.

**Figure 3:**
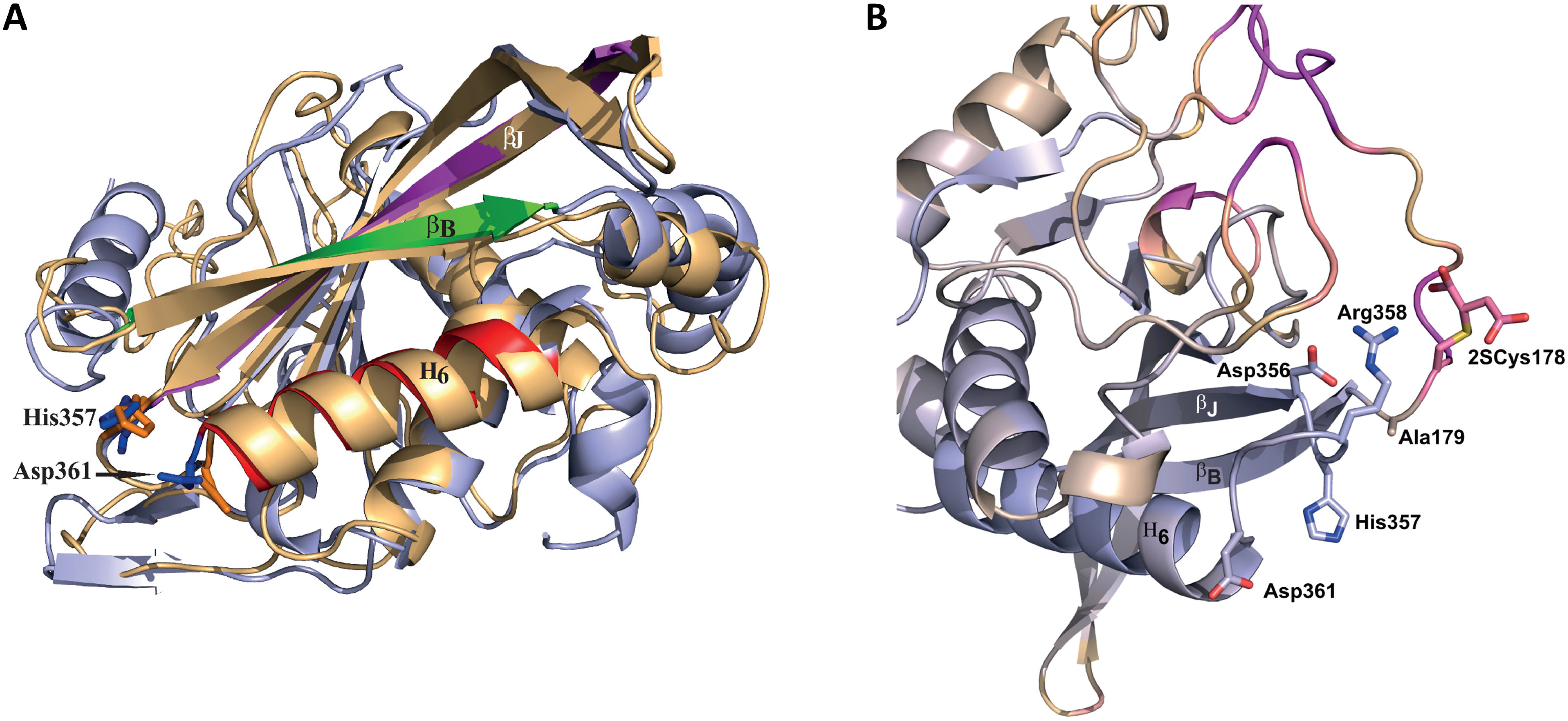
A, Superimposition of the succinated Cys178 (2SCys178) form of human DLST (hDLST) in orange and the unmodified hDLST in blue. Red, green, and purple colors in the unmodified form indicate the secondary structural elements that form the two substrate channels. B, The catalytic center of hDLST and its immediate spatial vicinity in the succinated (2SCys178) form, showing the degree of displacement relative to the unmodified structure. Blue color represents displacement <0.5 Å and purple indicates displacement >6.5 Å.

### The fumarate isomer maleate and DLST succinomimetics reduce the activity of the KGDH complex

We used maleate, the cis isomer of fumarate that has increased reactivity at physiological pH and is non-toxic compared to fumarate ester treatment (Bergholtz et al., 2020) to increase intracellular succination and determine if this impacted KGDHC activity. Treatment of cells with 5 mM maleate for 16 hrs led to a significant reduction in KGDHC activity (37.11% vs. untreated controls, p<0.01, Figure 4A), whereas 1 mM maleate did not decrease KGDHC activity. Reduced KGDHC activity with 5 mM maleate was paralleled by a marked elevation in protein succination (Figure 4B, 2SC panel), whereas less succination was detected with 1 mM maleate, but was still increased above controls. This exogenous addition of maleate differs from the endogenous production of fumarate observed in the Ndufs4 KO mice and could be expected to impact cytosolic proteins more than mitochondrial proteins. We examined the sources of ATP production in maleate treated cells and found that maleate caused a shift towards increased ATP synthesis linked to glycolysis (p<0.05 for 1 mM maleate vs. 0 mM maleate, and p<0.001 for 5 mM maleate vs. 0 mM maleate), and reduced ATP synthesis linked to mitochondrial function (p<0.0001 for both comparisons), but no changes in the total amount of ATP produced (Figure 4C) or cell survival. Total DLST levels were not affected by maleate treatment (Figure S4A, DLST panel), similar to our findings in the BS of the Ndufs4 KO mouse (Figure 1H). These data demonstrate specific reductions in KGDHC function in parallel with decreased mitochondrial function and a compensatory shift towards glycolysis during acute maleate induced succination. Separately, we also included the use of dimethyl fumarate (DMF), a reactive fumarate ester that results in protein succination (Piroli et al., 2019). Incubation of a KGDHC standard with DMF caused a dose-dependent decrease in the activity of the complex (Figure S4F). Furthermore, we confirmed decreased KGDHC activity in N1E-115 neurons incubated with DMF (Figure S4G). When we replaced DMF with dimethyl succinate, a DMF analog that does not modify cysteine residues, there was no loss of KGDHC activity (Figure S4G). Increased succination of a band at ∼48-50 kDa was observed in N1E-115 neurons after incubation with 50 and 100 µM DMF (Figure S4H, arrow in 2SC panel), and the band co-localized with DLST immunolabeling (Figure S4H, DLST panel). While fumarate esters are more reactive and are not comparable to physiological fumarate increases, they do demonstrate that elevated fumarate-derived succination contributes to reduced KGDHC activity.

**Figure 4:**
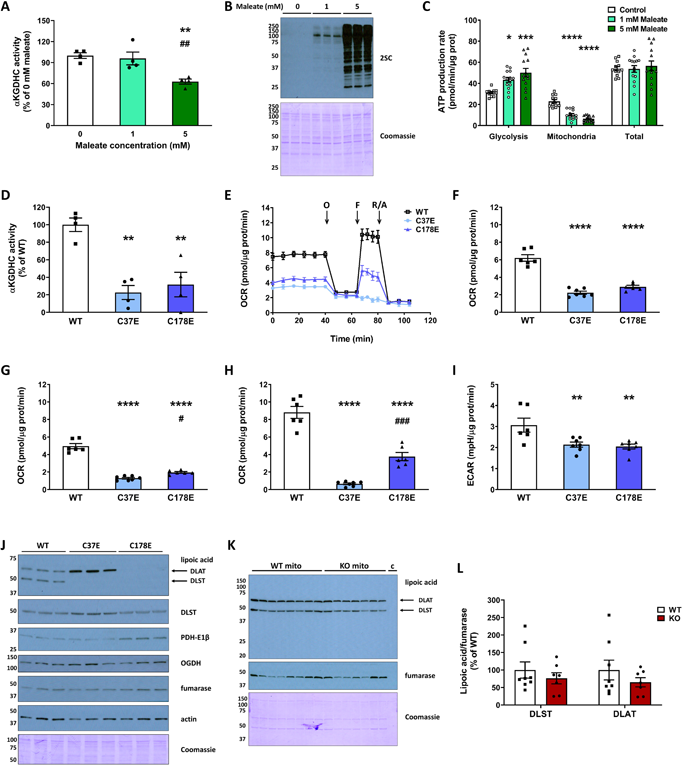
Maleate treatment and succinomimetic mutation of DLST reduce the activity of the KGDHC. *A-C*, KGDHC activity *(A)*, protein succination *(B)*, and ATP production rate linked to glycolysis and mitochondrial function *(C)* in DLST KO H838 cells expressing WT hDLST, after treatment for 16 h with 0-5 mM maleate. *D,* KGDHC activity in DLST KO H838 cells expressing WT hDLST or hDLST forms where Cys37 or Cys178 were mutated to Glu to mimic succination. *E*, Respiratory profile of the cells described in *(D)*. The oxygen consumption rate (OCR) was measured using a Seahorse XF analyzer, in basal conditions and after sequential injection of oligomycin (O), FCCP (F), and rotenone/antimycin A as described in Methods. *F-H*, Basal respiration (*F*), oxygen consumption coupled to ATP production (*G*), and maximal respiration (*H*), in the experiment described in (*E*). *I*, Extracellular acidification rate (ECAR) after oligomycin injection in the experiment described in (*E*). *J*, Detection of lipoic acid and mitochondrial markers (DLST, PDH-E1β, OGDH and fumarase) in the cells described in (*D*). Actin and Coomassie staining were used to verify even loading of the gel. *K*, Detection of lipoic acid moieties associated with DLST and DLAT (dihydrolipoyllysine-residue acetyltransferase, a subunit of the pyruvate dehydrogenase complex) in the BS of WT and Ndufs4 KO mice. In addition to the mitochondrial fractions (“mito”), a lane with a cytosolic fraction (“c”) was included as a negative control. *G,* Normalization of the signal intensity in *F*, relative to fumarase to account for mitochondrial protein levels. Normalization to Coomassie did not alter the lipoylation status (not shown). In panels *A*, *C*, *D*, and *F-I*, data expressed as mean ± SEM and comparisons by one-way ANOVA and *post hoc* Student-Neuman-Keuls test (*A* and *C,* *p<0.05, **p<0.01, ***p<0.001, and ****p<0.0001 for 1 mM and 5 mM maleate vs. 0 mM maleate, and ^##^ p<0.01 for 5 mM maleate vs. 1 mM maleate; *D*, and *F-I*, **p<0.01, and ****p<0.0001 for mutants vs. WT, and ^#^p<0.05 and ^###^p<0.001 for C178E vs. C37E). In panels *B* and *J*, molecular weight markers are shown on the left side.

We employed site-specific mutants to further study the impact of DLST succination on KGDHC functionality. H838 DLST-KO cells were previously generated by CRISPR-Cas9 technology (Allen et al., 2016, Remacha et al., 2019) and were transfected with a construct to express either wildtype (WT) hDLST, or with constructs where Cys37 or Cys178 were mutated to Glu to introduce a negative charge that mimics succination (the succinomimetics C37E and C178E). Glu is a larger amino acid than Asp and better mimics succination of cysteines (Kulkarni et al. 2019). We examined KGDHC activity in the cells expressing either WT hDLST, or the succinomimetic mutants C37E hDLST or C178E hDLST. As shown in Figure 4D, KGDHC activity was reduced in the mutant lines to 22.65 (C37E, p<0.01) and 30.58% (C178E, p<0.01) of the values obtained for the WT construct. In addition, the respiratory profile of the mutants was greatly affected (Figure 4E), with significant decreases in basal respiration (Figure 4F), respiration associated with ATP synthesis (Figure 4G), maximal respiration (Figure 4H), and the spare respiratory capacity (Figure S4B). No changes in respiration associated with proton leak or non-mitochondrial respiration (Figure S4C, D) were observed in the mutants. The extracellular acidification rate (ECAR) in basal conditions was not affected in the mutants (Figure S4E), but it was decreased after the addition of oligomycin (Figure 4I). This data confirms that the succinomimetic mutation of either cysteine in DLST profoundly impacts activity and the metabolic respiratory status of the cells.

Although we were unable to perform simulated molecular dynamics succination of Cys37, we observed that the C37E mutant was as effective in reducing KGDHC activity as the C178E mutant. We noted that Cys37 is only 6 residues away from the lipoylated lysine (Lys43^lipoyl^) in both mouse and human DLST [CEIETDK^lipoyl^]. We reasoned that succination of Cys37 in the Ndufs4 KO might interfere with DLST lipoylation, or that the reduced form of lipoate itself might be susceptible to succination. The production of succinyl-CoA by KGDHC employs the formation of a DLST-mediated succinyl-dihydrolipoate intermediate (Figure 1D), therefore we used an anti-lipoic acid antibody to investigate the lipoylation state of DLST. We first examined lipoylation in the cells expressing WT hDLST or the C37E and C178E succinomimetics as these would be expected to show a maximal effect. The antibody also detects another prominent lipoylated protein, dihydrolipoyllysine-residue acetyltransferase (DLAT), a lipoyl-containing subunit of the pyruvate dehydrogenase complex (PDH). The C178E mutation led to an almost complete loss of lipoic acid immunoreactivity in both DLST and DLAT (Figure 4J, lipoic acid panel), whereas the lipoylation state increased for DLAT and decreased for DLST for the C37E mutant (Figure 4J, lipoic acid panel). Total DLST levels were not affected in the C178E mutant (Figure 4J DLST panel) and appeared to be slightly increased in the C37E mutant (Figure 4J DLST panel). The levels of the PDH-E1β subunit increased slightly in the C178E mutant, perhaps in a compensatory manner, whereas levels of the OGDH subunit of KGDHC did not change markedly (Figure 4J). It is unclear why DLAT lipoylation was differentially impacted in the DLST-specific mutants, perhaps the loss of DLST activity promotes a separate oxidative lipoate modification precluding detection (Bailey et al. 2020., Vaubel et al., 2011). The pronounced reductions in DLST lipoylation do parallel the decreased KGDHC activity measured in these striking mutants. We next investigated if lipoylation was altered in the brain of the Ndufs4 KO mouse, however, we found that the lipoylated DLST and DLAT signal intensities were not reduced in the KO vs. WT BS (Figure 4K, L). Extensive attempts to detect peptides containing succinated Cys37 or succinated Lys43^lipoyl^ were also unsuccessful, although the difficulty in detecting lipoylated DLST modifications has previously been noted in ABHD11 deficient cells (Bailey 2020). The results of our simulation (Figure 3A, B) and the preserved lipoylation profile of DLST in the Ndufs4 KO brain further suggest that Cys178, and not Cys 37 or Lys43^lipoyl^, is the likely succination site in the Ndufs4 KO mouse.

### Defective DLST function and reduced KGDHC activity contribute to protein hyposuccinylation

We predicted that decreased synthesis of succinyl-CoA would contribute to reduced protein succinylation, an acyl CoA derived lysine modification that is distinct from cysteine succination (Zhang et al. 2011). Protein succinylation has been implicated in further modulating the activity of mitochondrial enzymes (Gibson et al., 2015; Park et al., 2013). DLST can act as a protein succinyl transferase (Gibson et al., 2015); therefore, the succination of DLST could contribute directly to decreased mitochondrial protein succinylation. This is outlined in Figure 5A, where the impact of increased fumarate-mediated generation of 2-succinocysteine (2SC) on DLST (DLST^2SC^) contributed to defective KGDHC activity, resulting in lower ATP via reduced SUCLA activity, decreased succinate levels and decreased lysine succinylation (Figure 5A). We investigated the levels of mitochondrial protein succinylation in DLST KO H838 cells expressing WT hDLST or the C37E and C178E mutants, since the mutants showed a striking reduction in KGDHC activity. The succinylation profile of mitochondrial proteins was notably decreased for the C178 mutant vs. WT across several select bands (Figure 5B), with a smaller reduction in succinylation in the C37E mutant vs. WT (Figure 5B). This suggests the C178E mutant resulted in a more significant deficit in succinyl-CoA levels. Cytosolic proteins were also hyposuccinylated in the C178E mutant (Figure S5C). We also examined the succinylation profile of mitochondrial proteins in DLST KO H838 cells expressing WT hDLST after 16 h treatment with 0-5 mM maleate. No significant change in succinylated proteins was detected (Figure S5A), similar results were obtained with N1E-115 neurons treated for 16 h with 0-100 μM DMF (Figure S5B), indicating that despite the reduction in KGDHC activity, this short treatment duration was not sufficient to impact the turnover of succinylated proteins.

**Figure 5:**
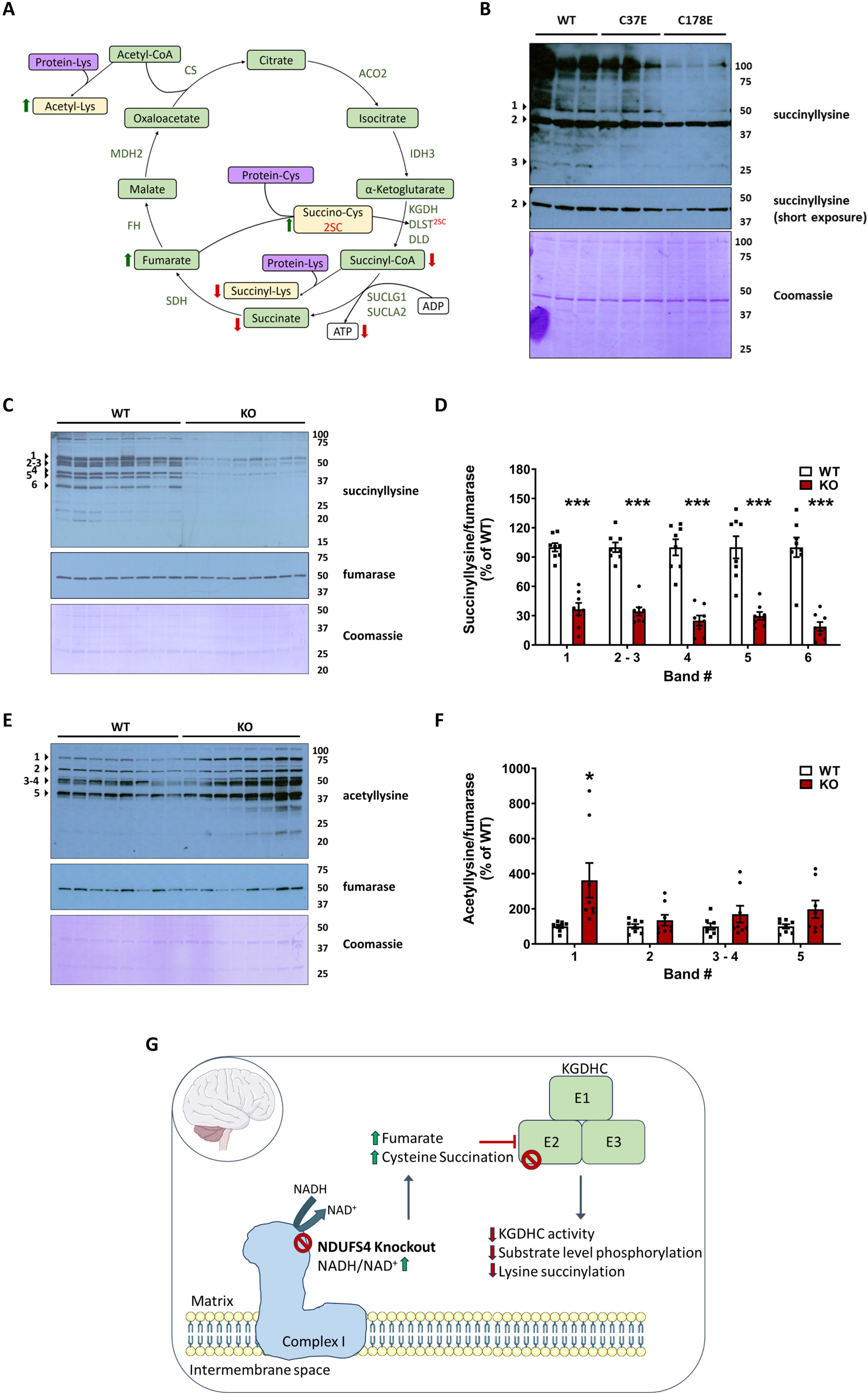
DLST succination and mutations that mimic DLST succination alter the levels of protein acylation *in vivo* and *in vitro*. *A*, Schematic illustrating TCA cycle deficits in the Ndufs4 KO mouse. The metabolic impact of increased fumarate derived succination of DLST (DLST^2SC^) contributes to defective KGDHC activity and results in lower ATP via reduced SUCLA activity, decreased lysine succinylation, and lower succinate levels in parallel with increased lysine acetylation. Relationships between metabolites (green boxes), protein lysine or cysteine residues (purple boxes) and protein modifications (yellow boxes) are indicated. *B,* Protein lysine succinylation in mitochondrial fractions obtained from DLST KO H838 cells expressing WT hDLST, C37E or C178E hDLST succinomimetic mutants., *C,* Protein lysine succinylation in mitochondrial fractions obtained from BS of WT and Ndufs4 KO mice. *D*, Quantification of prominent bands in *C* (arrowheads). *E*, Protein lysine acetylation in mitochondrial fractions obtained from BS of WT and Ndufs4 KO mice. *F*, Quantification of prominent bands in *E* (arrowheads). *G*, Overview of the TCA cycle perturbations occurring in the Ndufs4 knockout mouse brain. Reductive stress increases in parallel with fumarate and cysteine succination in the brainstem and olfactory bulb. Succination of DLST (E2 of KGDHC) lowers KGDHC activity and leads to reduced compensatory ATP production via substrate level phosphorylation. Mitochondrial hyposuccinylation is present due to decreased succinyl CoA production. These metabolic perturbations impacting the TCA cycle drive the chemical modification of proteins, extending the biochemical deficit beyond the loss of the Complex I component Ndufs4. In *B,* Coomassie Brilliant Blue staining (prominent band at 42 kDa) was used for the normalization of the succinyllysine signal. In *C* and *E*, re-probing to detect fumarase was used for mitochondrial protein normalization, followed by Coomassie Brilliant Blue staining. Molecular weight markers are shown on the right side. See also Figure S4. In *D* and *F*, results expressed as mean ± SEM were compared by unpaired t test (*p<0.05 and ***p<0.001 vs. WT).

We next investigated the mitochondrial protein succinylation profile in the brain of WT and Ndufs4 KO mice. Figure 5C demonstrates a remarkable global hyposuccinylation in the BS of the KO mice. Densitometric quantification of several prominent bands demonstrated >50% reduction in succinylation (Figure 5D). The Ndufs4 KO also showed decreased succinylation of cytosolic proteins in the BS, although less succinylation was detected in cytosolic vs. mitochondrial fractions (Figure S5D and S5E). Lysine succinylation changes were less evident in the nuclear protein fraction (Figure S5F, S5G), emphasizing that the greatest effect on succinylation status is within the mitochondria where fumarate is endogenously elevated in parallel with defective DLST function. Overall, reduced BS succinylation due to lower succinyl-CoA levels is further represented by decreased levels of succinate in the OB of the Ndufs4 KO (Figure S5H). Given that metabolite flux through KGDHC is impaired, we expected that acetyl CoA-derived mitochondrial lysine acetylation would be increased (Figure 5A), this is shown for select proteins in the Ndufs4 KO versus the WT (Figures 5E and 5F) and is aligned with the hyperacetylation recently described in a glutamatergic neuron specific Ndufs4 KO (Gella et al. 2020). Figure 5G summarizes the impact of increased reductive stress in the Ndufs4 KO, which is associated with brain region-specific increases in fumarate and protein succination. Cysteine succination of DLST (KGDHC E2) impairs the activity of the KGDHC, further compromising ATP generation via reduced substrate level phosphorylation and decreasing global mitochondrial lysine succinylation.

## Discussion

The results of the current study demonstrate that compromised DLST function due to cysteine succination by fumarate is sufficient to pronouncedly decrease KGDHC activity in the BS and OB of the Ndufs4 KO mouse, the two brain regions with the most profound pathological changes in this model of Leigh syndrome (Quintana et al., 2010). These important results link an OXPHOS genetic defect to impaired TCA function and defective succinyl-CoA production, a TCA intermediate necessary for SUCLA-mediated SLP (Kiss et al., 2013; Komlódi and Tretter, 2017) and succinylation reactions (see schematic in Figure 4F, Yang and Gibson, 2019). Our findings align with recent metabolomic efforts that suggest that defective glutamate/α-ketoglutarate metabolism distinguishes pathologically affected brain regions from unaffected regions (Johnson et al., 2020, Terburgh et al., 2021a). While redox changes modulate TCA cycle activity depending on the energy status of the cell, protein succination represents a static event that contributes to functional insufficiency of the affected proteins, elucidating an unrecognized pathological contributor in Complex I deficiency. The increase in fumarate levels despite significantly reduced KGDHC activity and succinate production likely reflect metabolic rewiring of the TCA cycle under conditions of reductive stress and metabolic acidosis, which limits the oxidative decarboxylation of malate (Nadtochiy et al., 2016) and allows both malate and fumarate accumulate (Figure 2A and S2D). In contrast, the oxidation of succinate is favored in the Ndufs4 KO to supply electrons for OXPHOS via FADH_2_ generation, supporting our observations of decreased succinate and increased fumarate and malate.

Previous reports on ATP synthesis levels in the Ndufs4 KO mouse vary depending on the tissue analyzed, with no differences in total cellular ATP content in immortalized fibroblasts (Valsecchi et al., 2012) and skeletal muscle (Alam et al., 2015; Kruse et al., 2008; Terburgh et al., 2019). However, neither fibroblasts nor skeletal muscle show overt pathology, nor did we find evidence of increased succination in the skeletal muscle of the Ndufs4 KO compared to WT mice (Piroli et al. 2016). Maximal mitochondrial ATP production in the whole brain is slightly decreased in Ndufs4 KO *versus* WT mice (Manjeri et al., 2016), and CI-linked ATP synthesis was also slightly decreased in isolated permeabilized neurons and astrocytes from mice with a spontaneous mutation leading to disruption of *NDUFS4* (Bird et al., 2014). We report that both total and SLP-linked ATP synthesis in mitochondria isolated from the OB of the Ndufs4 KO mouse were decreased when α-ketoglutarate, a preferential substrate for SLP, was used. A previous study using brain mitochondria from DLD^+/-^ and DLST^+/-^ mice showed diminished mitochondrial ATP efflux with fuel combinations that support SLP compared to WT mitochondria, and these mice also had a 20-48% decrease in KGDHC activity, which is consistent with our findings (Kiss et al., 2013). Our data provides novel mechanistic insight on the metabolic complexity of CI deficiency, elucidating how the mitochondria suffer from not only defective OXPHOS, but also have inadequate compensatory ATP generation via SLP. Reduced ATP production in the most susceptible neurons would accelerate their functional decline.

The role of KGDHC deficiency in neurodegeneration has been studied in the preclinical setting, including the genetic modulation of DLD and DLST; in both cases the KO mice were not viable (Johnson et al., 1997; Yang et al., 2009). DLST^+/-^ and DLD^+/-^ do not show overt symptoms but are susceptible to mitochondrial toxins used to replicate neurodegenerative diseases (Klivenyi et al., 2004; Yang et al., 2009). Pharmacological inhibition of KGDHC activity *in vitro* alters mitochondrial morphology and increases fission and mitophagy (Banerjee et al., 2016). Interestingly, the degree of KGDHC inhibition in the brains of Ndufs4 KO mice is aligned with that reported using pharmacological inhibitors of the complex to impair mitochondrial function (Banerjee et al., 2016). Similar to human OGDH mutation (Yap et al. 2020), human patients with mutations in the *LIPT1/2* genes that encode a lipoyltransferase necessary for DLST lipoylation develop Leigh Syndrome-like encephalopathies (Habarou et al., 2017; Stowe et al., 2018), supporting the important role for KGDHC deficiency in driving neuropathology. Brain regions from patients with Alzheimer’s disease (AD) show ∼57% reduced KGDHC activity (Butterworth and Besnard, 1990; Gibson et al., 1988). Of note, the expression of DLST in the human brain appears to be restricted exclusively to neurons (Dobolyi et al. 2020), supporting the notion that neuronal dysfunction is the triggering event for pathological lesion formation. Interestingly, DLST (E2) also serves as the E2 component of the 2-oxoadipate dehydrogenase complex (Nemeria et al., 2018), involved in lysine catabolism, and Ndufs4 KO metabolomics studies have shown evidence of lysine accumulation (Terburgh et al., 2021a), although dysregulated redox dynamics in the Ndufs4 KO brain may also contribute to impairments in this pathway.

Our current studies elucidate that mitochondrial stress driven by a genetic Complex I deficiency contributes to fumarate-driven reductions in KGDHC activity. Remarkably, neither of the two cysteines in mature DLST are part of the active site, but a recently determined structure demonstrated that Cys178 is in the spatial vicinity of the active site (Nagy et al., 2021), and Cys37 is proximal to the known lipoylation site. We employed both mathematical modeling and site-directed mutagenesis to further interrogate the impact of cysteine conversion to the larger, negatively charged succinocysteine. Modeling Cys178 succination demonstrated the formation of a new ionic interaction between the succinated cysteine and Arg358, displacing the normal Arg358-Asp356 salt bridge. Cys178 succination also altered the orientation of active site His357 and disrupted the H-bond that stabilizes His357 and the loop in which it resides and is predicted to result in less efficient coordination of the CoA and SLAM substrates and increased K_M_ values. These modeling observations strongly support the decreased succinyl CoA mediated succinylation profile and further succinate reductions observed in the Ndufs4 KO brainstem. While our studies focused on the brain regions affected by neuropathology, defective succinyl CoA production could also be anticipated to reduce global heme production, potentially contributing further to the iron dyshomeostasis recently described the Ndufs4 KO mouse (Kelly et al. 2023). We further employed a succinomimetic mutagenesis strategy to determine if switching cysteine to a larger, negatively charged amino acid (Glu) would alter KGDHC function. Remarkably, mutation of either cysteine had pronounced effects on KGDHC activity, mitochondrial respiratory capacity and DLST lipoylation status. Although the C37E mutation had the greatest impact on respiration, we noted that the striking hyposuccinylation profile observed in the C178E mutant was aligned with the hyposuccinylation observed in the Ndufs4 KO BS, and our modeling data suggested that altered DLST Cys178 succination might impair the ability of the active site to yield succinyl CoA. Since we were unable to define by mass spectrometry which DLST cysteine is succinated these data provide confidence that loss of either cysteine is sufficient to reduce the function of this central TCA cycle metabolic junction, Cys178 succination parallels metabolic defects observed in the Ndufs4 KO brainstem.

The profound mitochondrial hyposuccinylation profile detected has not been described in mitochondrial encephalopathies, and the additive functional impact of this succinylation decrease remains unknown in this model where a Complex I deficit already results in a lower ATP yield. KGDHC can succinylate proteins *in vitro* more efficiently than free succinyl-CoA, likely due to the succinyl-transferase activity of the E2 component (Gibson et al., 2015). In agreement with our observations in the Ndufs4 KO, KGDHC inhibitors reduce succinylation of cytosolic and mitochondrial proteins in primary neurons (Gibson et al., 2015). Mellid et al. have recently described hyposuccinylation due to DLST mutations in heritable human pheochromocytomas and paragangliomas (PPGL), noting marked decreases in succinylation of TCA cycle and glycolytic enzymes that may contribute to the tumorigenic capacity of these cells (Mellid et al. 2023). A recent report documented hypersuccinylation in *SUCLA2* deficiency, a mtDNA depletion syndrome (Gut et al. 2020). Hypersuccinylation contrasts with the hyposuccinylation that we report, but would be expected in a model where succinyl CoA is the accumulating metabolite versus the deficiency that we detect with reduced KGDHC function. It will be important to define which target proteins are differentially affected in these models as this may contribute to the heterogeneity of the molecular and clinical phenotype of these encephalopathies. While the *SUCLA2* insufficiency has some clinical similarity with *NDUFS4* deficiency, other features such as sensorineural deafness and mtDNA depletion differ from the CI defect. In contrast to cysteine succination, succinylation is reversible by specific sirtuin (SIRT) enzymes (Yang and Gibson, 2019). Overexpression of *SIRT5* increased survival and improved mitochondrial oxygen consumption in a zebrafish model of *SUCLA2* deficiency, but did not increase mtDNA content, suggesting that modulation of succinylation has functional effects on OXPHOS parameters (Gut et al. 2020).

Hypoxia has been shown to prolong lifespan and reduce brain lesions in the Ndufs4 KO mouse (Jain et al., 2016; Ferrari et al., 2017). Reduction in the oxygen content in the air normalizes the hyperoxia present in the brain of the Ndufs4 KO mouse to levels found in WT littermates breathing normal air (Jain et al., 2019). Interestingly, *in vitro* studies with N2a neural cells showed that hypoxia leads to increased mitochondrial protein succinylation (Chen et al., 2017), therefore it will be of interest to determine if hypoxia treatment can alleviate the decreased lysine succinylation observed in the Ndufs4 KO mouse. In addition, dimethyl α-ketoglutarate (DMKG), a cell permeable form of α-ketoglutarate, has been shown to delay neurological symptoms and increase the median lifespan of Ndufs4 KO by ∼40 days (Lee et al., 2019), and rapamycin treatment also pronouncedly increases α-ketoglutarate levels (Johnson et al. 2020) Overall increases in α-ketoglutarate would be expected to increase succinyl CoA and confer benefit by providing electrons to support Complex-II dependent respiration in neurons where ∼70% residual activity is present. Future targeted studies to reduce fumarate accumulation are expected to prevent the sustained damage caused by covalent protein succination during reductive stress, and improve the response to supplementary therapies, such as DMKG, to benefit neuronal survival during Leigh Syndrome.

## Conclusions

The biochemical mechanisms driving neuropathology in mitochondrial oxidative phosphorylation (OXPHOS) disorders, such as Leigh syndrome, are not well understood beyond the known ATP deficit. Therapeutic approaches to treat mitochondrial diseases employ vitamins and antioxidants, yet these do not significantly slow the rate of neurological decline. In contrast, reductive stress is associated with metabolic acidosis and altered tricarboxylic acid (TCA) cycle metabolism. We demonstrate that fumarate levels are increased in brain regions of the *NDUFS4* knockout mouse model of mitochondrial Complex I deficiency. Increased fumarate reacts covalently with protein thiols to generate the chemical modification 2-succinocysteine (protein succination). We report that the specific succination of dihydrolipoyllysine-residue succinyltransferase (DLST) irreversibly reduces the activity of the α-ketoglutarate dehydrogenase complex (KGDHC), confirmed by modeling cysteine succination and succinomimetic DLST mutagenesis. This persistent deficit worsens the mitochondrial OXPHOS derived ATP deficit by limiting compensatory substrate level phosphorylation (SLP) via succinyl CoA ligase. Defective KGDHC activity also reduces succinyl CoA mediated lysine succinylation, a distinct acyl modification, in the affected brain regions of the Ndufs4 knockout mouse. The data mechanistically demonstrates how altered TCA cycle metabolism drives the chemical modification of proteins to disrupt metabolic homeostasis; and provides a novel biochemical basis for neuronal metabolic decline during Complex I OXPHOS deficiency.

## Author Contributions

G.G.P and N.F. designed the research. G.G.P., R.S.M., A.M.M., H.H.S., W.E.C, M.D.W., O.O., and S.M. performed experiments. G.G.P., R.S.M., A.M.M., H.H.S., W.E.C, M.D.W., O.O., A.A. and N.F. analyzed data. A.C. provided new reagents. G.G.P., A.M.M., and N.F. wrote the paper.

## Acknowledgements

This work was supported by the National Institutes of Health (R01NS126851, R56NS116174, R01NS092938, to N.F), F31DK108559 (to A.M.M)), National Science Foundation (1828059) and a University of South Carolina Research Foundation ASPIRE-I awards (to N.F and G.G.P). This study was supported by the Hungarian Scientific Research Fund (OTKA grant 143627, to A.A.) and the Ministry of Innovation and Technology of Hungary (TKP2021-EGA-25 grant, to A.A. (project no. TKP2021-EGA-25 has been implemented with the support provided by the Ministry of Innovation and Technology of Hungary from the National Research, Development, and Innovation Fund, financed under the TKP2021-EGA funding scheme)). Additional support from project PI22/01490 (to A.C) from the Instituto de Salud Carlos III (ISCIII) through the “Acción Estratégica en Salud” (AES), cofounded by the European Regional Development Fund (ERDF).

## MATERIALS AND METHODS

### RESOURCES TABLE

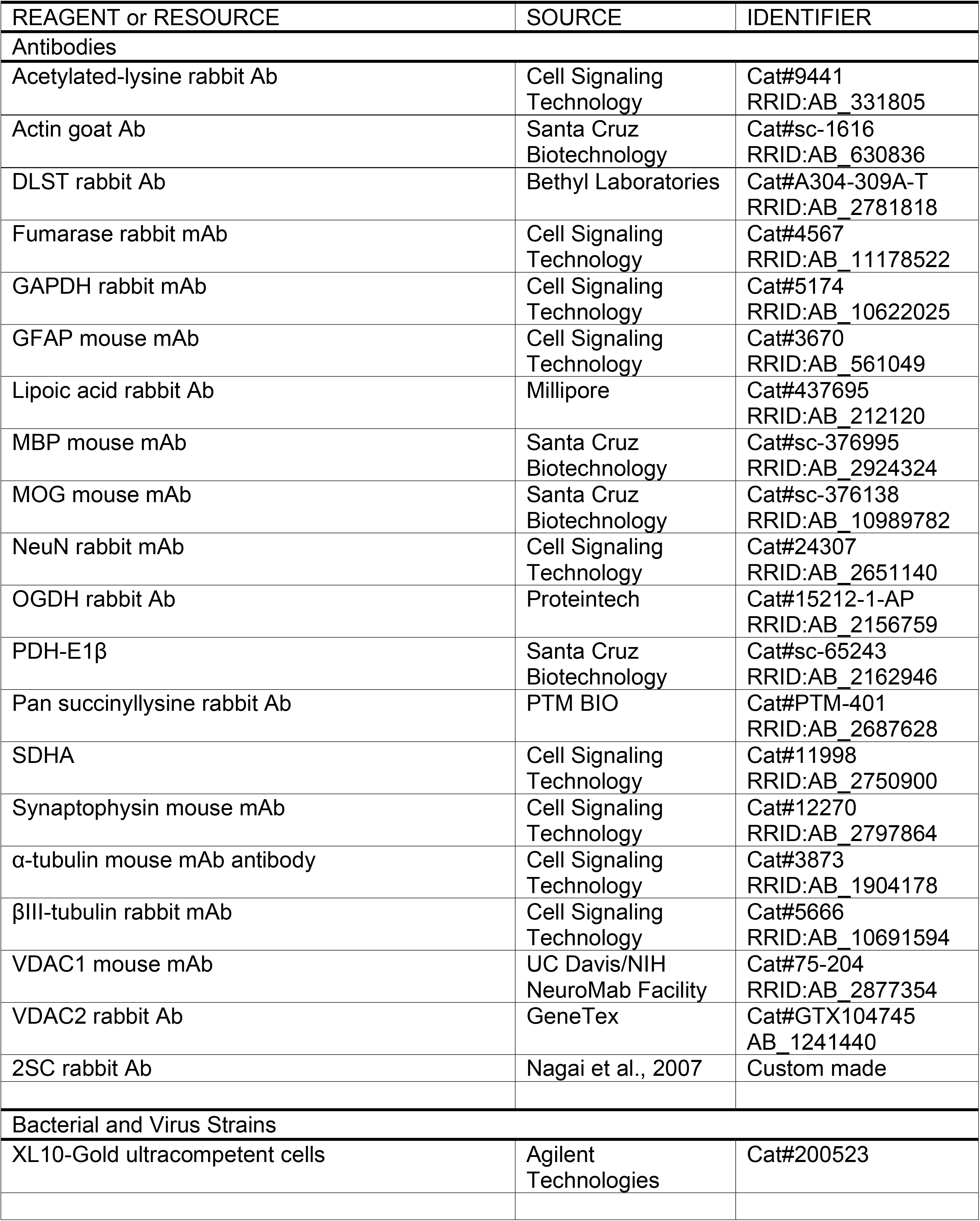

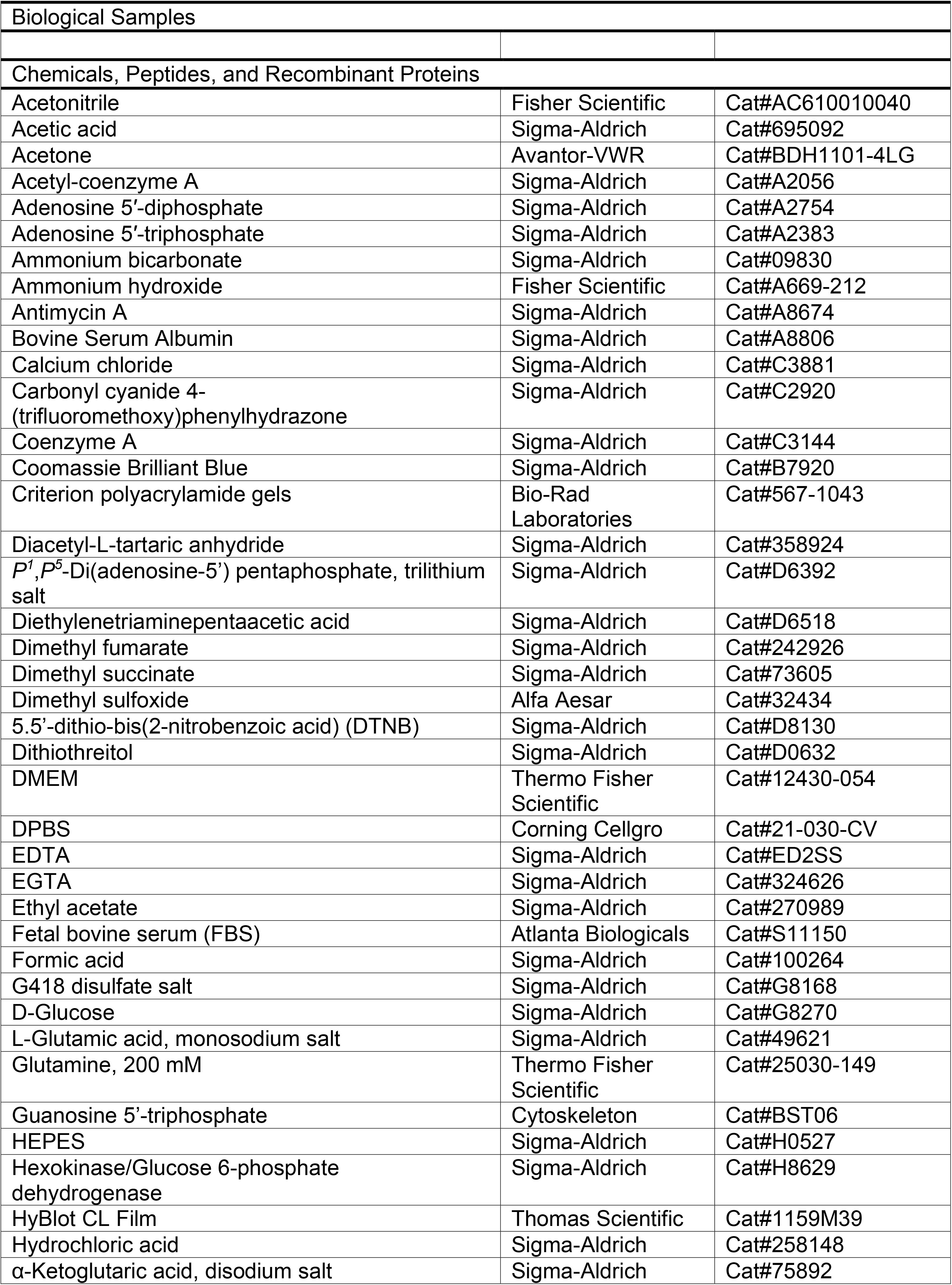

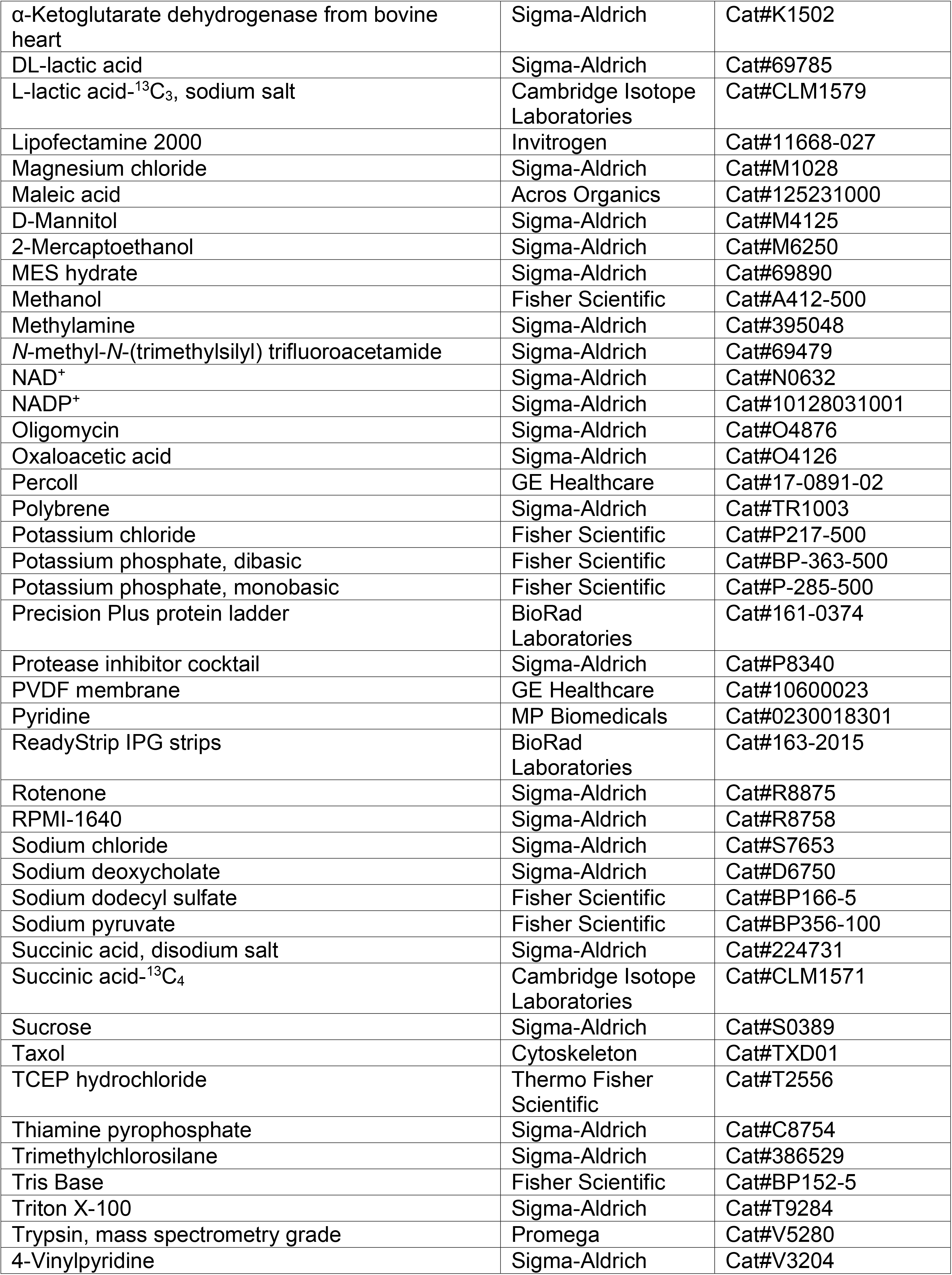

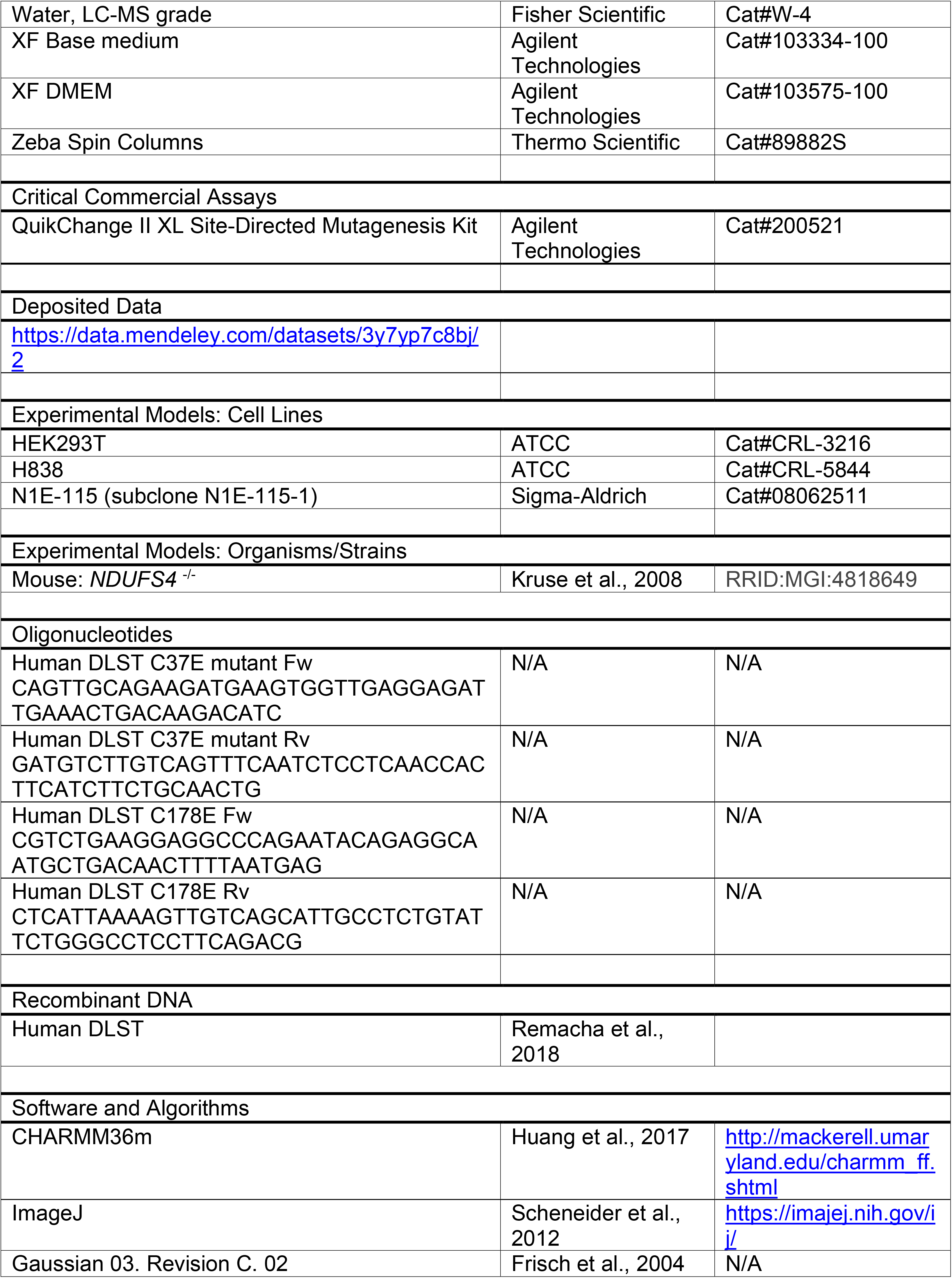

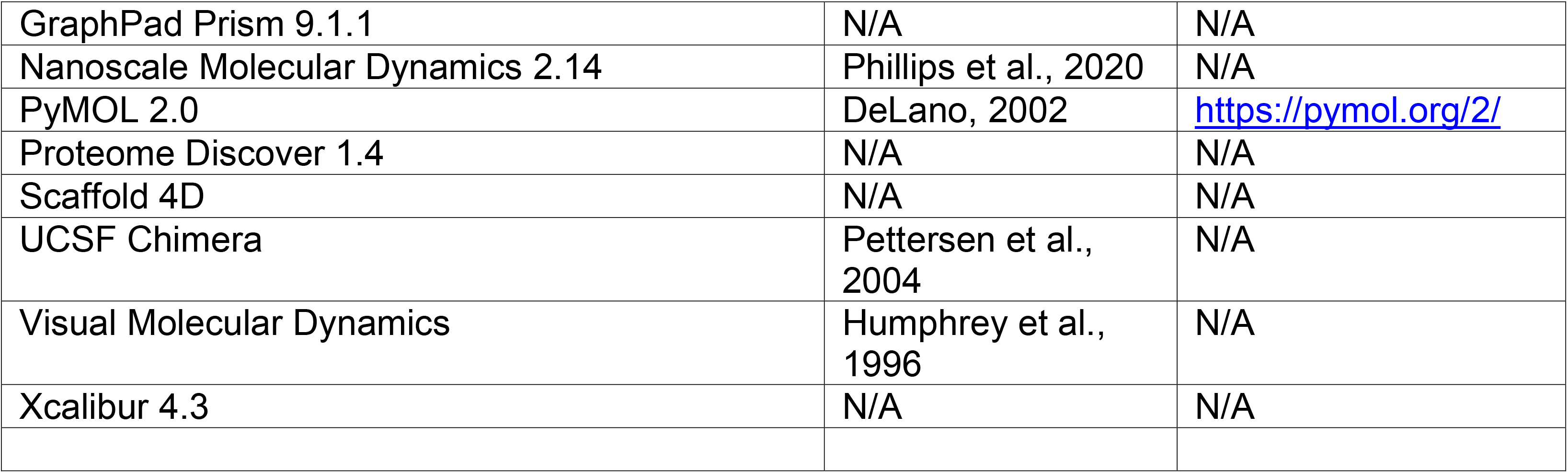

### EXPERIMENTAL MODELS

#### Mice

Animal care and use procedures were carried out in accordance with the National Institutes of Health Guide for the Care and Use of Laboratory Animals and approved by The University of South Carolina Animal Care and Use Committee. Ndufs4^+/-^ mice (129/Sv x C57BL/6 mixed background), originally obtained from Dr. Richard Palmiter (University of Washington, Seattle, WA) (Kruse et al., 2008; Quintana et al., 2010), were mated in trios (one male and two females), switched to a breeder’s rodent diet (LabDiet 5058) and weaned at 21 days of age. Weaned male and female mice were housed in a pathogen-free animal facility under a 12h light/dark cycle (lights on from 7AM to 7PM) and received Teklad 8904 rodent diet and water ad libitum. Genotyping was performed as previously described (Piroli et al., 2016), and both male and female mice were used. Ndufs4^+/+^ and Ndufs4^-/-^ mice were sacrificed by decapitation under deep isoflurane anesthesia at 8 weeks of age. The brain was removed from the skull, and the brainstem (BS), olfactory bulbs (OB) and cerebellum were dissected. Other tissues including liver were also collected. For some protocols fresh brain regions were used immediately; on other occasions the tissues were snap frozen in liquid nitrogen and stored at -80 °C until further use (see METHODS DETAILS).

#### N1E-115 Cells

N1E-115 neuroblastoma cells (subclone N1E-115-1) were expanded in non-differentiation medium containing 90% DMEM and 10% FBS, as previously described (Piroli et al., 2016; Piroli et al., 2019). At 80% confluence, the cells were differentiated into neurons in the presence of 2% FBS and 1.25% dimethyl sulfoxide in DMEM for 5 days.

#### H838 Cells

DLST-KO was generated in H838 human lung epithelial adenocarcinoma cells by CRISPR-Cas9 technology (Allen et al., 2016). The vectors used for the expression of DLST mutants C37E and C178E were obtained through mutagenesis of a previously generated *DLST* cDNA construct (Remacha et al., 2018) using the QuikChange II XL Site-Directed Mutagenesis Kit and the primers described in the Key Resources Table above. Lentivirus-based constructs (for both the Cys mutants and the WT hDLST) were made according to the standard protocol of the RNAi Consortium from the Broad Institute (http://portals.broadinstitue.org/gpp/public/resources/protocols). Briefly, HEK293T cells were transfected with a lentiviral vector mix including the corresponding *DLST* cDNA construct and Lipofectamine 2000 (ratio DNA:Lipofectamine was 1:2.5). Media was replaced 16 h after transfection. The virus-containing supernatant was collected 24 h later and added to the H838 DLST-KO cells together with 8 mg/mL of Polybrene. This step was repeated the following day for a second infection. Drug selection started 12 h later, with RPMI-1640 medium containing 10% FBS and 400 μg/ml G418. Selection proceeded until all uninfected control cells were dead. The cells were then switched to RPMI-1640 medium containing 10% FBS and 200 μg/ml G418.

### METHODS

#### Mouse brain homogenates for immunoblotting

Cleared homogenates were prepared from frozen brain regions in radioimmunoprecipitation assay (RIPA) buffer (50 mM Tris-HCl pH 7.4, with the addition of 150 mM NaCl, 1 mM EDTA, 0.1% Triton X-100, 0.1% SDS, 0.5% sodium deoxycholate, 2 mM diethylenetriaminepentaacetic acid and a protease inhibitor cocktail). Homogenization was performed by pulse sonication at 2 watts using a Model 100 sonic dismembrator (Fisher Scientific, Fair Lawn, NJ) for 30 s prior to resting on ice for 30 min. The homogenate was clarified from nuclei and unbroken cells by centrifugation at 900 x *g* for 10 min at 4°C. Protein in the supernatants was precipitated with 9 volumes of cold acetone for 10 min on ice. After centrifugation at 3,000 x *g* for 10 min and removal of the acetone, the protein pellet was resuspended in RIPA buffer. These cleared homogenate fractions were used for detection of succinated proteins (Figures 1A, 1G and S1D).

Considering the abundance of tubulin in total brain homogenates, the high level of tubulin succination, and the location of the ∼48-50 kDa succinated band that we describe in this manuscript immediately below tubulin, we followed three different protocols to enrich the ∼48-50 kDa band and at the same time deplete them of tubulin.

*First*, we subjected mouse BS samples to an *in vitro* tubulin polymerization protocol that we previously described (Piroli et al., 2014). Briefly, frozen mouse BS samples were reduced to a powder with a pestle in a mortar containing liquid nitrogen. The pulverized tissue was then resuspended in cold Mes/glutamate buffer (0.1 M Mes, pH 6.8, containing 0.5 mM MgCl_2_, 1 mM EGTA, 1 M glutamate, 1 mM DTT and a protease inhibitor cocktail), in a volume ratio of 1:1.5 (powder:buffer). The suspension was pulse sonicated for 5 intervals of 10 s. The total protein homogenate was then centrifuged at 30,000 x *g* at 4°C for 15 min to generate a pellet (P1) that was analyzed by immunoblotting (Figure 1B), and the supernatant (S1) was subjected to microtubule polymerization by addition of 20 μM taxol and 1 mM GTP, followed by incubation for 30 min at 37°C. Following further centrifugation at 30,000 x *g* for 30 min at 37°C, the supernatant of microtubules (SM) was removed, desalted through Zeba Spin Columns (MW cut-off 7 kDa, Thermo Scientific) and analyzed by immunoblotting (Figure S1B); the microtubular pellet (MP) composed of >95% tubulin was not used.

Second, we prepared gliosomes and synaptosomes from BS and cerebellum samples according to Carney, et al. (2014), with minor modifications. Briefly, fresh BS and cerebellum samples were homogenized in 0.32 M sucrose, 1 mM EDTA, 10 mM HEPES pH 7.4 using a glass-teflon homogenizer; the resulting crude homogenates were centrifuged at 1,000 x *g* at 4°C for 5 min. The supernatants (cleared homogenates) were then loaded on top of a gradient composed of layers containing 3, 7, 10 and 20% Percoll in the homogenization buffer; a fraction of the cleared homogenate was saved for immunoblotting comparative purposes. The gradients were centrifuged at 33,500 x *g* at 4°C for 6 min without brakes, and the interfaces between 3 and 7% Percoll (gliosomes), 10 and 20% (synaptosomes) and the loose pellet at the bottom of the tubes were collected and washed with the homogenization buffer to remove the Percoll. All the collected fractions and the cleared homogenates were analyzed by immunoblotting (Figures 1C and S1C).

Third, we prepared purified mitochondrial fractions according to Kayser, et al. (2016), with minor changes. Briefly, fresh BS samples were homogenized in 225 mM mannitol, 75 mM sucrose, 1 mM EGTA, 5 mM HEPES pH 7.2 containing 1 mg/ml fatty acid-free BSA and a protease inhibitor cocktail using a glass-teflon homogenizer; the resulting crude homogenates were centrifuged at 1,100 x *g* at 4°C for 2 min. The supernatants (cleared homogenates) were added to Percoll to a 5% concentration, and then layered on top of 15% Percoll; a fraction of the cleared homogenate was saved for immunoblotting comparative purposes. The gradients were centrifuged at 18,500 x *g* at 4°C for 10 min; the top layer, the interface and the lower layer were removed and the loose pellets at the bottom of the tubes were collected and washed with 250 mM sucrose, 0.1 mM EGTA, 5 mM HEPES pH 7.2 to remove the Percoll. The mitochondrial pellets and the cleared homogenate were analyzed by immunoblotting for succinated proteins (Figure 1F).

#### Cell Treatments to Increase Protein Succination

N1E-115 cells were treated during the final 16 h of differentiation with 0-100 µM dimethyl fumarate (DMF) prepared in Dulbecco’s PBS (DPBS) as a 100X stock and passed through a 0.22 μm pore nitrocellulose filter for sterility; some of the wells received dimethyl succinate (DMS) in the same range of concentrations as a negative control. H838 DLST-KO cells receiving a construct to restore WT hDLST were treated during 16 h with 0-5 mM maleate prepared in DPBS and filtered as described above. The cell collection and subcellular fractionation is described in detail in the section “Measurement of the KGDHC activity in purified bovine heart complex and cells”.

#### Subcellular Fractionation of Cells for Immunobloting

The subcellular fractionation was performed according to Manuel, et al. (2020), with minor modifications. H838-DLST KO cells expressing the hDLST WT and cysteine mutants were rinsed twice with PBS and then with homogenization buffer (10 mM Tris-HCl, pH 7.4, containing 0.32 M sucrose, 1 mM EDTA, 1mM dithiothreitol and a protease inhibitor cocktail). Homogenization buffer was added and cells were gently scraped and homogenized in glass-glass homogenizers, followed by six syringe passes through a 25 G needle. This initial homogenate was centrifuged at 600 x *g* for 10 min at 4°C in a microcentrifuge, and the supernatant was further centrifuged at 18,000 x *g* for 20 min in a microcentrifuge to isolate mitochondrial enriched pellets; the supernatant was saved as cytosol. These fractions were used for immunoblotting (Figures 3J, 4A, and S4C)

#### One-dimensional PAGE and Western Blotting

Western blotting was performed as described previously (Piroli et al., 2014; Piroli et al., 2016; Piroli et al., 2019). Ten to fifty µg of proteins were run on 12% gels (Criterion, BioRad) and transferred to PVDF membranes. Immunoblotting was performed with antibodies listed in the Key Resources Table. The preparation of the polyclonal anti-2SC antibody has been described previously (Nagai et al., 2007). In most cases, membranes were stripped with 62.5 mM Tris, pH 6.8, containing 2% SDS and 0.7% 2-mercaptoethanol for 15 min at 65°C prior to reprobing. Chemiluminescent signals were captured on photographic film (HyBlot CL).

#### Two-dimensional Gel Electrophoresis and Western Blotting

Isoelectric focusing on pI 4-7, 11 cm strips and two-dimensional (2D) gel electrophoresis was performed in an Ettan IPGphor II device (Amersham Biosciences) as described previously (Piroli et al., 2014; Piroli et al., 2016). The 2D gels were transferred onto PVDF membranes to detect protein succination by western blotting; the blots were then stripped and subjected to DLST detection.

#### Measurement of the α-ketoglutarate dehydrogenase complex activity in brain regions

The activity of the α-ketoglutarate dehydrogenase complex (KGDHC) was measured by following the formation of NADH at 340 nm according to Yang, et al. (2009), with minor modifications. BS and OB frozen samples were homogenized in 50 mM Tris-HCl pH 7.2 containing 1 mM dithiothreitol, 0.2 mM EGTA, 0.4% Triton X-100 and a protease inhibitor cocktail using a glass-teflon homogenizer. The crude homogenates were centrifuged at 2,000 x *g* for 4 min at 4°C; the supernatants were used to determine KGDHC activity. One hundred µl of a reaction mix containing 50 mM Tris-HCl pH 7.8, 1 mM MgCl_2_, 1 mM CaCl_2_, 0.5 mM EDTA, 0.3 mM thiamine pyrophosphate, 0.1% Triton X-100 and 1 mM dithiothreitol were added to the wells of a 96-well plate, followed by 50 μl of an assay mix containing 50 mM Tris-HCl pH 7.8, 3 mM NAD^+^ and 0.75 mM coenzyme A, and then 25 µl of the 2,000 x *g* supernatants (see above) containing 5-10 µg protein (or homogenization buffer for the blanks). A baseline reading every min for 10 min with the temperature set at 30°C was taken at 340 nm in a plate reader (Tecan Safire^2^, TECAN Systems Inc.). After the addition of 25 μl 3 mM α-ketoglutarate in 50 mM Tris-HCl pH 7.8 (or 50 mM Tris-HCl pH 7.8 for “no substrate” controls), further readings were taken for an additional period of 60 min. The readings in the absence of α-ketoglutarate were subtracted from those in the presence of the substrate, and these corrected values were plotted as a function of time to calculate the V_max_ from the slopes. Each sample was run in triplicates that were averaged; the n size was 3-4 for each experimental group. The results were converted to mU/mg of protein considering E_NAD(340)_ = 6.22 cm^-1^mM^-1^ and a path length of 0.4 cm.

#### Measurement of the KGDHC activity in purified bovine heart complex and cells

*In vitro* succination effects on the activity of a commercially available bovine heart KGDHC standard was studied after incubation for 16 h at room temperature with a range of 0-2 mM DMF following the protocol described above for brain regions; dithiothreitol was replaced with 300 µM tris(2-carboxyethyl) phosphine (TCEP) to avoid the possible reaction of DMF with dithiothreitol. The results of the KGDHC activity were expressed as percentage of the control group (no DMF added). The KGDHC activity was also measured in N1E-115 cells treated with DMF or DMS and H838 cells treated with maleate (see “Cell Treatments” above) with minor modifications. Briefly, the medium was removed; the cells were rinsed 3 times with DPBS and gently scraped with 50 mM Tris-HCl pH 7.2 containing 1 mM dithiothreitol, 0.2 mM EGTA, 0.4% Triton X-100 and a protease inhibitor cocktail. The extracts were homogenized in a glass-glass homogenizer, the resulting homogenates were centrifuged at 2,000 x *g* for 4 min at 4°C, and the supernatants were used to determine KGDHC activity following the protocol described above for brain regions. The results of the KGDHC activity were expressed as percentage of the control group (no DMF, DMS or maleate added).

#### Measurement of citrate synthase activity in brain regions

The activity of citrate synthase was measured according to Leek, et al. (2001), with minor modifications. Briefly, frozen brain regions were homogenized in 100 mM monobasic/dibasic potassium phosphate pH 7.4, containing 5 mM EDTA, 5 mM EGTA and a protease inhibitor cocktail with a glass-teflon homogenizer (ratio buffer to tissue was 20:1). The initial homogenate was sonicated twice for 20 s to disrupt the mitochondrial membrane, and then centrifuged at 2,000 x *g* for 4 min at 4°C. The resulting supernatants were then frozen, thawed, and subjected to a new cycle of sonication, freezing and thawing, with a final dilution 1:20 in the homogenization buffer before proceeding to the assay. The measurement was performed by mixing 180 μl of 100 mM Tris pH 8.1, containing 0.1 mM 5.5’-dithio-bis(2-nitrobenzoic acid) (DTNB), 0.2 mM acetyl-coenzyme A and 0.5 mM oxaloacetic acid with 20 μl of the samples (3-6 μg protein); blank wells did not contain oxaloacetic acid. The side reaction between coenzyme A and DTNB, with formation of a mercaptide ion, was followed at 30°C in a plate reader at 412 nm at 20 s intervals for 20 cycles. The readings in the absence of oxaloacetic acid were subtracted from those in the presence of the substrate, and these corrected values were plotted as a function of time to calculate the V_max_ from the slopes. Each sample was run in triplicates that were averaged; the n size was 4 for each experimental group. The results were converted to mU/mg of protein considering E_mercaptide ion(412)_ = 13.6 cm^-1^mM^-1^ and a path length of 0.4 cm.

#### Molecular simulation of Cys178 succination effects on DLST functionality

Molecular dynamics simulations were performed as described previously (Ambrus et al., 2015, Nagy et al., 2021), with adaptations as described below. Molecular graphics software Visual Molecular Dynamics (VMD; Humphrey et al., 1996) and UCSF Chimera (Pettersen et al., 2004) were used to model the structure of both the unmodified human DLST (hDLST) and the succinated Cys178 variant; the 6H05 PDB cryo-EM structure was used as a starting point. hDLST trimer coordinate and protein structure files were then generated according to the symmetry operators as well as the atom types and connectivity definitions of the CHARMM36m force field (Huang et al., 2017). The effects of succination on Cys178 were characterized *via* running an *ab initio* calculation using the Gaussian 03W software package (Frisch et al., 2004) to carry out a density functional theory modeling at the B3LYP/6-31G^++^(d,p) level of theory. The obtained results together with the CHARMM36m force field data of cysteine and glutamic acid were used to set the parameters − force constants, equilibrium bond lengths and angles, partial charges, van der Waals parameters − of the succinated cysteine residue.

Preliminary results indicated that the region before Asn161 has very high flexibility that led to unstable dynamics simulation. This prompted us to model only residues 161-386. The trimeric protein was embedded in an ionized (K^+^ and Cl^-^, at 150 mM concentration) water sphere, applying the TIP3P water model. The system was set up in a way that the ionization states of the amino acid side chains mimicked the pH∼7 condition, with the δ-N of the histidine imidazole ring being protonated. The program Nanoscale Molecular Dynamics (NAMD; Phillips et al., 2020) was used to run a 50 ns long simulation of the unmodified and succinated hDLST. Energy minimization of the constructed protein was carried out using the conjugate gradient method in 200,000 cycles. The subsequent molecular dynamic simulation run was carried out under Linux using the CUDA-enabled 2.14 version of the NAMD software package. The simulation parameters were set up as follows: (i) in terms of time steps, 1 fs intervals were applied, (ii) calculation of electrostatic interactions (in this spatially limited system) was performed by the particle mesh Ewald method (Herce et al., 2007) applying a real-space cut-off distance of 12 Å and grid width of 1 Å, (iii) switching distance for non-bonded electrostatics and van der Waals interactions was 10 Å and the scaled 1-4 exclusion scheme was applied, and (iv) final temperature of the system was set to 310.15 K. During the pre-equilibration run for gaining a microcanonical ensemble, the temperature was augmented from 10.15 K to the final temperature in 10 K increments; 2,000 simulation steps were performed at each temperature, except for the final one, where an additional 190,000 steps were also carried out. The main run was controlled *via* Langevin dynamics under isobaric-isothermal condition; the temperature and pressure were maintained at 310.15 K and 101,325 Pa (1 atm; Langevin piston Nose-Hoover method included in the NAMD package; Phillips et al., 2020; Rudin et al., 2013), respectively. The trajectory was sampled between 35.7 and 36.7 ns (stable region highlighted in Figure S2J) and the average coordinates, calculated by using the VMD smooth trajectory script, were energy minimized in 20,000 cycles. The final succinated structure was analyzed *via* superimposing to the unmodified monomeric/dimeric/trimeric structure. Interatomic distances and root-mean-square deviations (RMSD) were computed by UCSF Chimera. Structure visualization was performed by the program Pymol.

#### Substrate level phosphorylation measurements

Mitochondrial isolation and measurement of ATP synthesis was performed according to Komlódi and Tretter (2017), with minor modifications. Freshly obtained BS and OB from Ndufs4 KO mice and WT littermates were immediately homogenized in 5 mM Tris-HCl pH 7.4 containing 225 mM mannitol, 75 mM sucrose and 1 mM EGTA using a glass-teflon homogenizer. The initial homogenates were centrifuged at 1,300 x *g* for 3 min, and the pellets were resuspended, washed twice in the homogenization buffer and saved as a nuclear fraction (see below). The supernatants of the 1,300 x *g* centrifugation were further centrifuged at 20,000 x *g* for 10 min. The resultant supernatant was saved as cytosol (see below), and the pellet was resuspended in 15% Percoll and then layered on top of a discontinuous Percoll gradient consisting of 23% and 40% layers. The gradients were then centrifuged at 30,700 x *g* for 8 min using no brake at the end. Immunoblotting experiments determined that the interface between the 23% and 40% Percoll and the loose pellet in the 40% Percoll layer were enriched in mitochondrial markers and devoid of myelin (Figure S2G). Consequently, these two fractions were pooled together, topped with homogenization buffer and centrifuged for 10 min at 16,600 x *g*; and the pellet was re-suspended in homogenization buffer and subsequently centrifuged again at 6,300 x *g* for 10 min. The resulting purified mitochondrial pellet was re-suspended in 5 mM Tris-HCl pH 7.4 containing 225 mM mannitol and 75 mM sucrose, and immediately used for mitochondrial ATP synthesis measurements. The assay medium consisted of 20 mM HEPES pH 7.0 containing 0.1 mM EGTA, 125 mM KCl, 2 mM K_2_HPO_4_, 1 MgCl_2_, and 0.025% fatty acid-free bovine serum albumin, with the addition of 3 mM NADP^+^, 5 mM glucose, 300 µM AP5 (P^1^,P^5^-Di(adenosine-5’) pentaphosphate; an inhibitor of adenylate kinase), 0.75 U/ml hexokinase, and 0.25 U/ml glucose-6-phosphate dehydrogenase, in a final volume of 200 µl. ATP formation was detected through the phosphorylation of glucose by added hexokinase, coupled to the oxidation of the resulting glucose 6-phosphate by added glucose 6-phosphate dehydrogenase, with formation of NADPH. NADPH synthesis was followed at 340 nm in a plate reader as described for the measurement of α-ketoglutarate dehydrogenase complex activity (see above). Basal measurements started upon addition of the mitochondrial samples (5 µg/well) and ADP (2 mM) for 10 min at 37°C, followed by the addition of α-ketoglutarate or succinate (both at 5 mM); some wells also received oligomycin (8 µM) to determine total ATP synthesis (oxidative phosphorylation + substrate level phosphorylation) in the absence of oligomycin, and substrate level phosphorylation in the presence of oligomycin. Corrections in the absence of substrate were applied, and these corrected values were plotted as a function of time to calculate the ATP synthesis from the slopes. Each sample was run in triplicates that were averaged; the n size was 7-12 for each experimental group. The results were converted to nmoles ATP/min/mg of protein using an ATP curve (range: 0-20 nmoles). The purified mitochondrial fractions were also used for immunoblotting experiments (succinylation in Figure 4C, acetylation in Figure 4E, succination in Figure S2H, and detection of protein lipoylation in Figure S3F). The nuclear fractions obtained after the centrifugation at 1,300 x *g* were used for immunoblotting experiments (succinylation in Figure S4F), whereas the cytosols obtained after the centrifugation at 20,000 x *g* were used for immunoblotting experiments (succinylation in Figure S4D).

#### Metabolic Measurements with the Seahorse Extracellular Flux Analyzer

H838 cells were seeded on V7 Seahorse cell culture microplates at a density of 30,000 cells/well. After 2 days in culture, two different experiments were performed with these cells.

*Experiment 1* was a mitochondrial stress assay designed to test the effect of DLST cysteine mutations on mitochondrial functionality. The cells were switched to XF Basal Medium supplemented with 10 mM glucose and 2 mM glutamine (same concentrations as in the RPMI medium used to grow the cells), and the oxygen consumption rate (OCR) was measured with a Seahorse XFe24 Extracellular Flux Analyzer in basal conditions (no drugs injected), and after injections of oligomycin (2 µM), carbonyl cyanide 4-(trifluoromethoxy)phenylhydrazone (FCCP, 0.25 µM), and rotenone/antimycin A (3 µM/4 µM) to determine ATP production, proton leak, maximal respiration, spare respiratory capacity and non-mitochondrial respiration, as previously described (Piroli et al., 2019).

*Experiment 2* was an ATP production rate assay designed to test the contribution of glycolysis and mitochondria to the total ATP synthesis in cells expressing WT hDLST when treated for 16 h with 0-5 mM maleate to increase protein succination. The cells were switched to XF DMEM supplemented with 10 mM glucose, 2 mM glutamine and 1 mM sodium pyruvate. The extracellular acidification rate (ECAR) and the OCR were then measured in basal conditions and after sequential injections of oligomycin (1.5 µM), and rotenone/antimycin A (0.5 µM for both drugs) to determine the relative contribution of glycolysis and mitochondrial metabolism to ATP synthesis, and the total amount of ATP produced.

For both experiments, the wells washed 3 times with cold PBS after completion of the assay and the plates were stored at -70°C prior to the measurement of the total protein content to normalize individual measurements.

#### Determination of Protein Content

The protein content in all the brain and cell homogenates, including samples for immunoblotting and those for the determinations of enzymatic activity was determined by the method of Lowry, et al. (1951).

#### Metabolite Quantification

TCA cycle metabolite quantification was performed in an adaptation of previous methods (Oldham et al. 2016, Manuel et al. 2020). Metabolites were extracted by adding 1ml ice-cold 80% methanol to snap-frozen olfactory bulbs. The tissue was dissociated in pre-cooled glass-teflon homogenizers and transferred to 1.5 ml Eppendorf tubes. Homogenized samples were briefly pulse-sonicated and frozen for 15 min at -70°C. After thawing, the samples were centrifuged at 15,000 x *g* for 10 min at 4°C. The supernatants were transferred to new Eppendorf tubes and the extracts were dried by centrifugal evaporation. The protein pellet was resuspended in RIPA buffer for quantification by the Lowry method.

The quantification of L-2-hydroxyglutarate by LC-MS was performed following chiral derivatization with diacetyl-L-tartaric anhydride (DATAN). 50 μM L-2-hydroxyglutarate-^13^C_5_ internal standard was added to tissues prior to extraction, and to all racemic standards. Dried tissue metabolite extracts and standards were resuspended in 50 μl DATAN prepared in acetonitrile:acetic acid (4:1, v/v) to 50 mg/ml and heated to 70 °C for 2 h. The samples were cooled to room temperature and centrifuged at 1000 x *g* for 5 min. The derivatized samples were further diluted to 100 μl with acetonitrile, vortexed, and transferred to LC vials for analysis. LC separation was performed on Thermo Vanquish Flex liquid chromatograph using a PEEK coated SeQuant ZIC HILIC column (Merck Millipore). The column was 2.1 mm x 150 mm with 3.5 μm particles and was preceded by an equivalent 20 mm long guard column. One μl of sample was separated by isocratic elution with 15% mobile phase A (10% 200 mM formic acid:90% LC-MS grade water) and 85% mobile phase B (10% 200 mM formic acid:90% acetonitrile) at a total flow rate of 200 μl/min. The formic acid was titrated to pH 3.25 with ammonium hydroxide. Negative ion electrospray mass spectra were acquired on a Thermo Q-Exactive HF-X quadrupole-Orbitrap in MS/MS mode with the Orbitrap resolution set at 7,500. Precursor ions were isolated by the quadrupoles and fragmented in the HCD cell at 10 ev. The product ions were mass analyzed in the Orbitrap. XCalibur 4.3 software was used to construct chromatograms of the transitions 363>147 (L-2-HG) and 368>152 (L-2-HG-^13^C_5_). The peak areas were normalized to the areas of the ^13^C internal standard and the analyte mass was normalized to the tissue weights.

The quantification of TCA cycle intermediates was performed by GC-MS at the David H. Murdock Research Institute (DHMRI, Kannapolis, NC). Prior to derivatization the extracts were resuspended in ethyl acetate and transferred to GC-MS vials. The samples were dried with N_2_, and an internal standard (100 μM succinate-^13^C_4_) was added to each of the samples and standards. The samples and standards were derivatized with 200 μl of methylamine (20 mg/ml in pyridine) at 30°C for 90 min, followed by drying under N_2_. This was followed by the addition of 120 μl of N-methyl-N-(trimethylsilyl) trifluoroacetamide (MSTFA) with 1% trimethylchlorosilane (TMCS); the mixture was incubated at 70°C for 60 min. The derivatized product was stored in a - 20°C freezer for one hour prior to GC/MS analysis. An Agilent 7890A GC system, coupled to an Agilent 5975C electron ionization (EI) mass selective detector (MSD) was used to analyze the TMS-derivatized samples. Select ion monitoring (SIM) was performed and the peak areas obtained were normalized to the internal standard. Absolute quantitation for all metabolites was performed based on standard curves obtained from the normalized reference standards, and the final metabolite concentrations were normalized to tissue weights or total protein content.

#### Protein identification by liquid chromatography-tandem mass spectrometry (LC-MS/MS)

The in-gel protein digestion method used was previously described (Piroli et al., 2016; Piroli et al., 2019). Briefly, samples of the pellets obtained from the synaptosomal-enriched preparations (20 µg of protein) were resolved by SDS-PAGE in 7.5% gels; the gels were stained with Coomassie Brilliant Blue and the bands of interest in the ∼48-50 kDa region were excised. The gel pieces were destained, washed with 50 mM ammonium bicarbonate in 50% acetonitrile, and dehydrated with 100% acetonitrile; proteins were then reduced with 10 mM dithiothreitol and alkylated with 170 mM 4-vinylpyridine. Protein digestion was carried out overnight at 37°C in the presence of 500 ng sequencing grade modified trypsin in 50 mM ammonium bicarbonate. After gel extraction with 5% formic acid in 50% acetonitrile, the samples were analyzed in a blinded manner on a Dionex Ultimate 3000-LC system (Thermo Scientific) coupled to a Velos Pro Orbitrap mass spectrometer (Thermo Scientific). The LC solvents were 2 % acetonitrile/0.1 % formic acid (Solvent A) and 80 % acetonitrile/0.1 % formic acid (Solvent B); the water used for these solvents was LC-MS grade. Peptides were first trapped on a 2 cm Acclaim PepMap-100 column (Thermo Scientific) with Solvent A at 3 µl/min. At 4 min the trap column was placed in line with the analytical column, a 75 μm C18 stationary-phase LC PicoChip Nanospray column (New Objective). The peptides were eluted with a gradient from 98%A:2%B to 40%A:60%B over 30 min, followed by a 5 min ramp to 10%A:90%B that was held for 10 min. The Orbitrap was operated in data-dependent acquisition MS/MS analysis mode and excluded all ions below 200 counts. Following a survey scan (MS1), up to 8 precursor ions were selected for MS/MS analysis. All spectra were obtained in the Orbitrap at 7500 resolution. The DDA data were analyzed using Proteome Discover 1.4 software with SEQUEST algorithm against the uniprot_ref_mouse database (2014-10-03 version, 52,474 proteins) with XCorr validation >2 (+2) or >2.5 (+3). An allowance was made for 2 missed cleavages following trypsin digestion. No fixed modifications were considered. The variable modifications of methionine oxidation (M^OX^), proline hydroxylation (P^OX^), cysteine pyridylethylation (C^PE^, 105.058) or cysteine succination by fumarate (C^2SC^, 116.011) were considered with a mass tolerance of 15 ppm for precursor ions and a mass tolerance of 10 ppm for fragment ions. The results were filtered with a false discovery rate of 0.01. For all DLST peptides identified the spectra were manually inspected to confirm the abundance of product ions. The software Scaffold 4D was used to generate the spectrum in Figure 1E.

### QUANTIFICATION AND STATISTICAL ANALYSIS

Results were expressed as means ± standard error of the mean (SEM). For the enzymatic activities, a description of the calculations is presented in the corresponding subheading under METHOD DETAILS. Statistical comparisons between two experimental groups were performed using unpaired *t* tests; when three or more groups were compared, one-way ANOVA followed by Student-Neuman-Keuls’ test was applied. The n size for each particular experiment is shown in the Figures and represents the number of individual biological replicates. Differences were considered statistically significant when a *P* value < 0.05 was achieved. The software Image J (NIH) was used for the quantification of band intensities by densitometry (Tanis et al., 2015). The software GraphPad Prism V9.1.1 was used for statistical analysis and graphs.

## DATA AVAILABILITY

The mass spectrometry. RAW files used to identify dihydrolipoyllysine-residue succinyltransferase peptides have been deposited in Mendeley Data with the identifier: https://data.mendeley.com/datasets/3y7yp7c8bj/2

## Supplementary Figure Legends

**Figure S1**: Supplementary Figure 1.

*A*, Formation of 2-(S-succino)cysteine (2SC) following covalent modification of protein cysteine residues in protein by fumarate. *B*, Detection of a distinct succinated band at ∼48-50 kDa (arrow) in a supernatant obtained after high speed centrifugation of *in vitro* polymerized microtubules from Ndufs4 KO BS homogenates. Tubulin and Coomassie panels show even loading of proteins (30 µg prot/lane). *C*, The cerebellum (CB) also shows evidence of neuropathology in the KO mice. Similar to the BS (see Figure 1B), a band at ∼48-50 kDa shows increased succination after synapotosomal- and gliosomal-enriched preparation of CB of Ndufs4 KO mice (arrow in 2SC panel). The pellets are enriched in the mitochondrial marker VDAC2. Coomassie panel shows even protein load (15 µg prot/lane) within each fraction group. *D*, Similar to BS (see Figure 1E), a train of succinated spots is observed in the region of ∼48-50 kDa and pH 5.5-6.3 (rectangular area) in Ndufs4 KO mice CB homogenates. *E*, MS/MS peptide identification for dihydrolipoyllisine-residue succinyltransferase component of 2-oxoglutarate dehydrogenase complex. Raw data were searched with the Sequest HT node of Proteome Discover 1.4 (SP1), using the uniprot_ref_mouse database. Variable modifications of methionine oxidation (M^OX^), proline oxidation (P^OX^), cysteine pyridylethylation (C^PE^), and cysteine succination by fumarate (C^2SC^) were considered. The peptides identified are reported from 3 individual KO mice analyzed. *F*, Total protein homogenates of brainstem, liver and olfactory bulb were compared to demonstrate increased succination in brain, particularly on select protein in the KO vs. WT. Immunodetection of OGDH, SDHA, fumarase, citrate synthase and actin was performed to demonstrate no loss of TCA cycle protein components in the KO.

In panels *B-D*, molecular weight markers are shown on the left side.

**Figure S2**: Supplementary Figure 2.

*A-D*, Pyruvate (*A*), citrate (*B*), oxaloacetate (*C*), and malate (*D*) contents in the OB of Ndufs4 KO and WT mice. *E-F,* Citrate synthase activity in the OB (*E*) and BS (*F*) of WT and Ndufs4 KO mice. *G*, Purity of the fractions obtained for substrate level phosphorylation (SLP) measurement (See Method Details section). The initial homogenate (H), cytosol (C), and different fractions obtained after separation of the initial particulate pellet through a Percoll density gradient, including layer 1 (L1), layer 2 (L2), interface 2 (I2) and layer 3 (L3) were separated by SDS-PAGE. After protein transfer, mitochondrial markers (SDHa and VDAC1), cytosolic markers (α-tubulin) and myelin markers (MOG and MBP) were detected by immunoblot. Fractions I2 and L3 were combined for the SLP assay due to enrichment of mitochondria and lack of cytosolic components and myelin. *H*, Confirmation that succination of a band at ∼48-50 kDa in mitochondrial fractions from BS used for SLP measurements is limited to Ndufs4 KO mice.

In panels *A-F*, results expressed as mean ± SEM, and comparisons performed by unpaired t test (* p<0.05 and ** p<0.01 for Ndufs4KO vs. WT). In panels *G* and *H*, molecular weight markers shown on the left side.

**Figure S3**: Supplementary Figure 3.

*A*, Comparison of human and mouse DLST sequences. Cysteine (C) residues present in the mature mitochondrial isoform are highlighted in blue, and the site of lipoylation (K) is highlighted in red. Residues underlined in green are important for the 3D structure of DLST in terms of functionality, and their interactions are greatly affected if Cys178 is succinated (see C and *D* in this figure, and Figure 3A and B). *B*, Structural motions in the hDLST trimer (blue) and its backbone (orange) in the course of the molecular dynamic simulation, relative to the starting conformation. *C*, Representative interatomic distances between residues having roles in the stability and functionality of the hDLST active site when Cys 178 is unmodified or succinated. Specific atoms in groups as follows: ^a^: NH2, ^b^: OD2, ^c^: NH1, ^d^: SG, ^e^: NH2, ^f^: OE1, ^g^: OD1, ^h^: NH. Minimum distances were measured in a single monomer, with measurements in the other two monomers always showing <16% difference. *D*, Dynamics of the Ala179-Asp356 distance in the succinated (Cys178) hDLST trimer. The average distance is 4.38 Å. *E*, Residue displacements in the trimeric succinated (Cys178) hDLST relative to the trimeric unmodified structure.

In panels *D* and *E*, each chain in the trimer is designated by a different color: blue, orange, and grey are for chains A, B, and C, respectively.

**Figure S4**: Supplementary Figure 4.

*A*, Total DLST levels in DLST KO H838 cells expressing WT hDLST treated for 16 h with maleate (0-5 mM) to cause protein succination. Actin and Coomassie staining were used to verify even loading. *B-D*, Spare respiratory capacity (*B*), proton-leak associated respiration (*C*), and non-mitochondrial respiration (*D*) in DLST KO H838 cells expressing WT hDLST or hDLST forms where Cys37 or Cys178 were mutated to Glu to mimic succination (C37E, C178E). *E*, Basal extracellular acidification rate (ECAR) in the same cell groups described in *B-D*. *F*, Effect of the incubation with dimethyl fumarate (DMF) for 16 h on the activity of a bovine heart KGDHC standard. *G*, Effect of the addition for 16 h of DMF or dimethyl succinate (DMS) to differentiated N1E-115 neurons on the KGDHC activity. *H*, Protein succination in N1E-115 neurons exposed to DMF. The succinated band at ∼48-50 kDa (arrow) co-localized with DLST immunoreactivity.

Results expressed as mean ± SEM. In panels *B-E*, comparisons by one-way ANOVA and *post hoc* Student-Neuman-Keuls test (** p<0.01, and **** p<0.0001 for mutants vs. WT, and ^###^ p<0.001 for C178E vs. C37E). In panel *G*, comparisons by unpaired t test (NS). In panels *H* and *I*, comparisons by one-way ANOVA and *post hoc* Student-Neuman-Keuls test (** p<0.01, and *** p<0.001 for treated vs. control, and ^##^ p<0.01 for 2 mM DMF vs. 0.5 mM DMF). In panels *A* and *I*, molecular weight markers shown on the left side.

**Figure S5**: Supplementary Figure 5.

*A*, Protein lysine succinylation in mitochondrial fractions obtained from DLST KO H838 cells expressing WT hDLST, after treatment for 16 h ± 5 mM maleate. *B*, Protein lysine succinylation in mitochondrial fractions obtained from differentiated N1E-115 neurons, after treatment for 16 h of 0-100 μM dimethyl fumarate (DMF). *C,* Protein lysine succinylation in cytosolic fractions obtained from DLST KO H838 cells expressing WT hDLST, C37E or C178E hDLST succinomimetics. *D*, Protein lysine succinylation in cytosolic fractions obtained from BS of WT and Ndufs4 KO mice. *E*, Quantification of prominent bands in *D* (arrowheads), relative to actin. *F*, Protein lysine succinylation in nuclear fractions obtained from BS of WT and Ndufs4 KO mice. *G,* Quantification of prominent bands in *F* (arrowheads), relative to NeuN. *H,* Reduced production of succinate in Ndufs4 KO vs. WT controls.

In *A-D* and *F,* molecular weight markers are shown on the right, and prominent succinylated bands are shown on the left side of the panels. Coomassie Brilliant Blue staining shows equal loading of the lanes.

In *E, G* and *H*, results expressed as mean ± SEM were compared by unpaired t test (*p<0.05, ***p<0.001 and ****p<0.0001 vs. WT).

## Notes

http://dx.doi.org/10.17632/3y7yp7c8bj.2

### Competing Interest Statement

The authors have declared no competing interest.

### Summary of Updates

Protein simulation studies and mutant studies have been performed and added.

