## Supplementary figures and images for "Defective Function of α-Ketoglutarate Dehydrogenase Exacerbates Mitochondrial ATP Deficits during Complex I Deficiency"

### Supplementary Figure 1

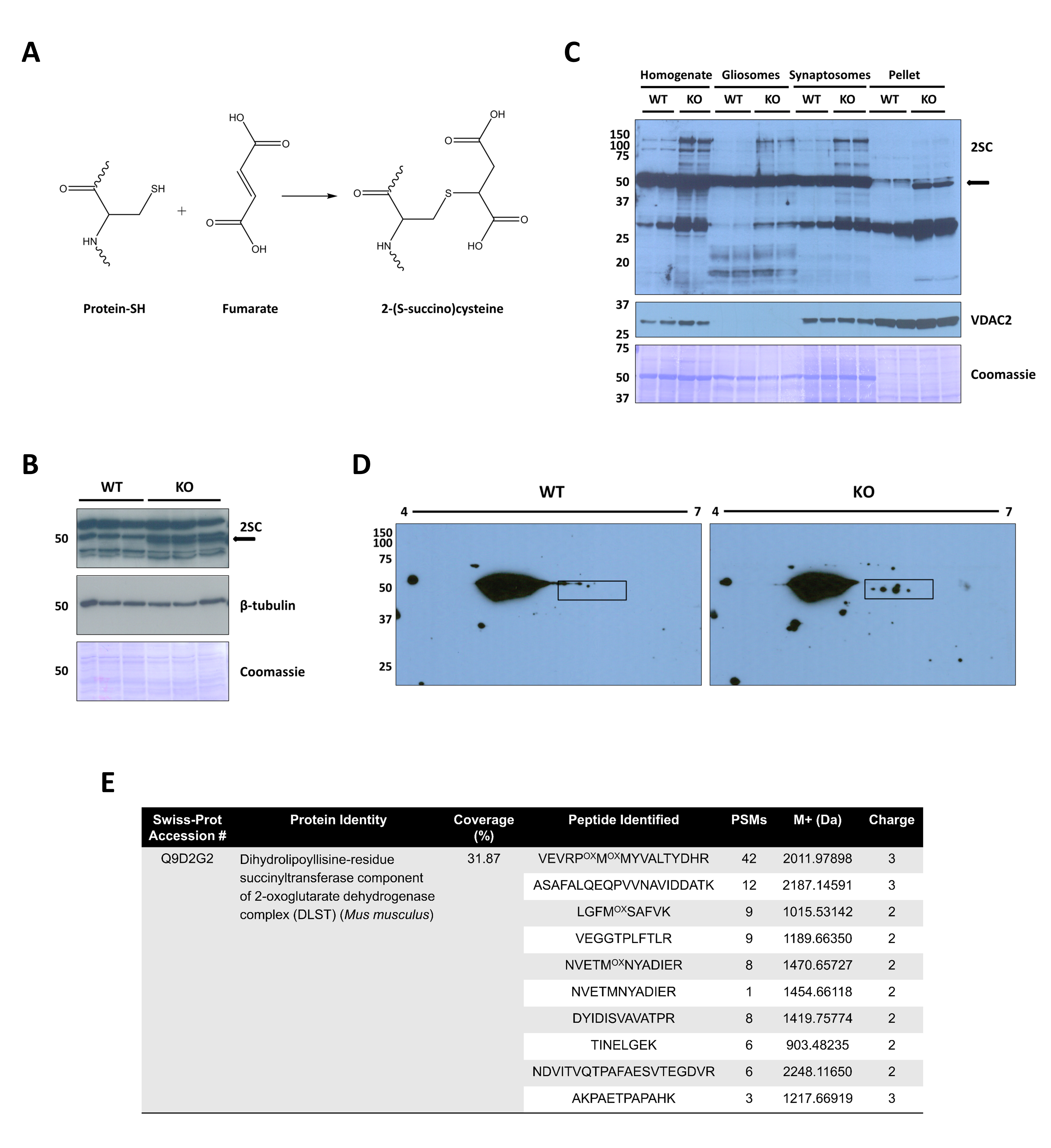

### Supplementary Figure 2

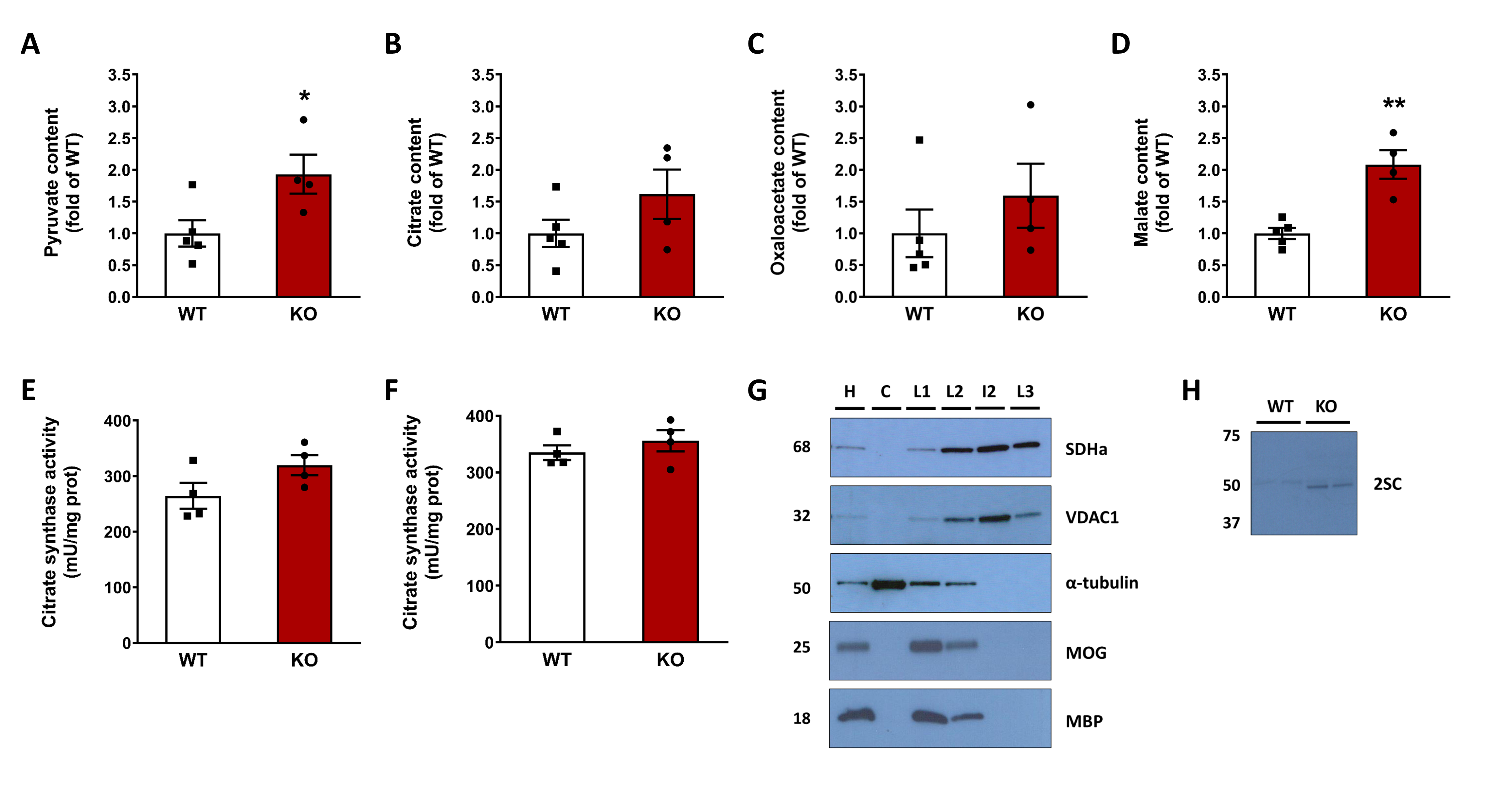

### Supplementary Figure 3

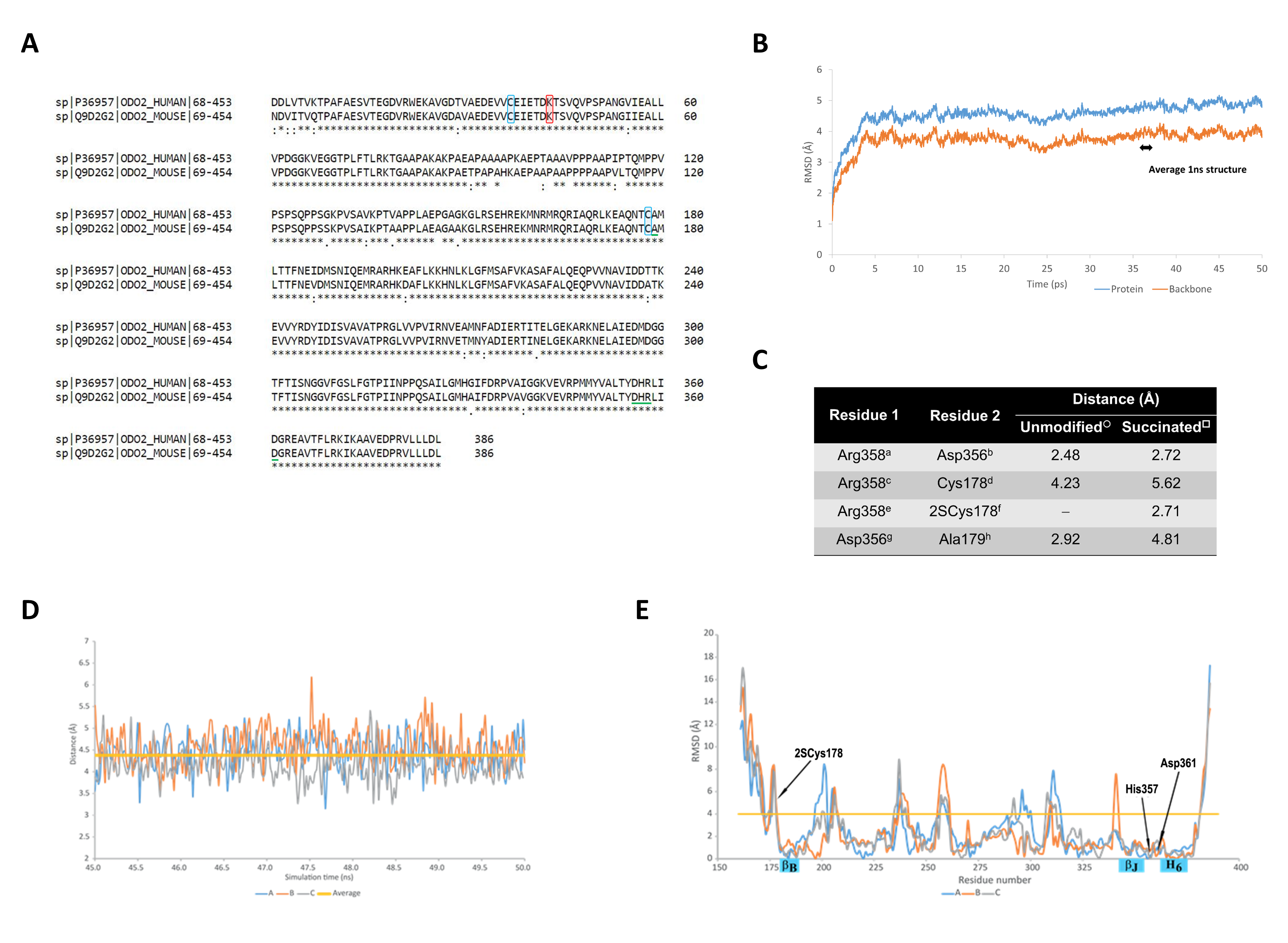

### Supplementary Figure 4

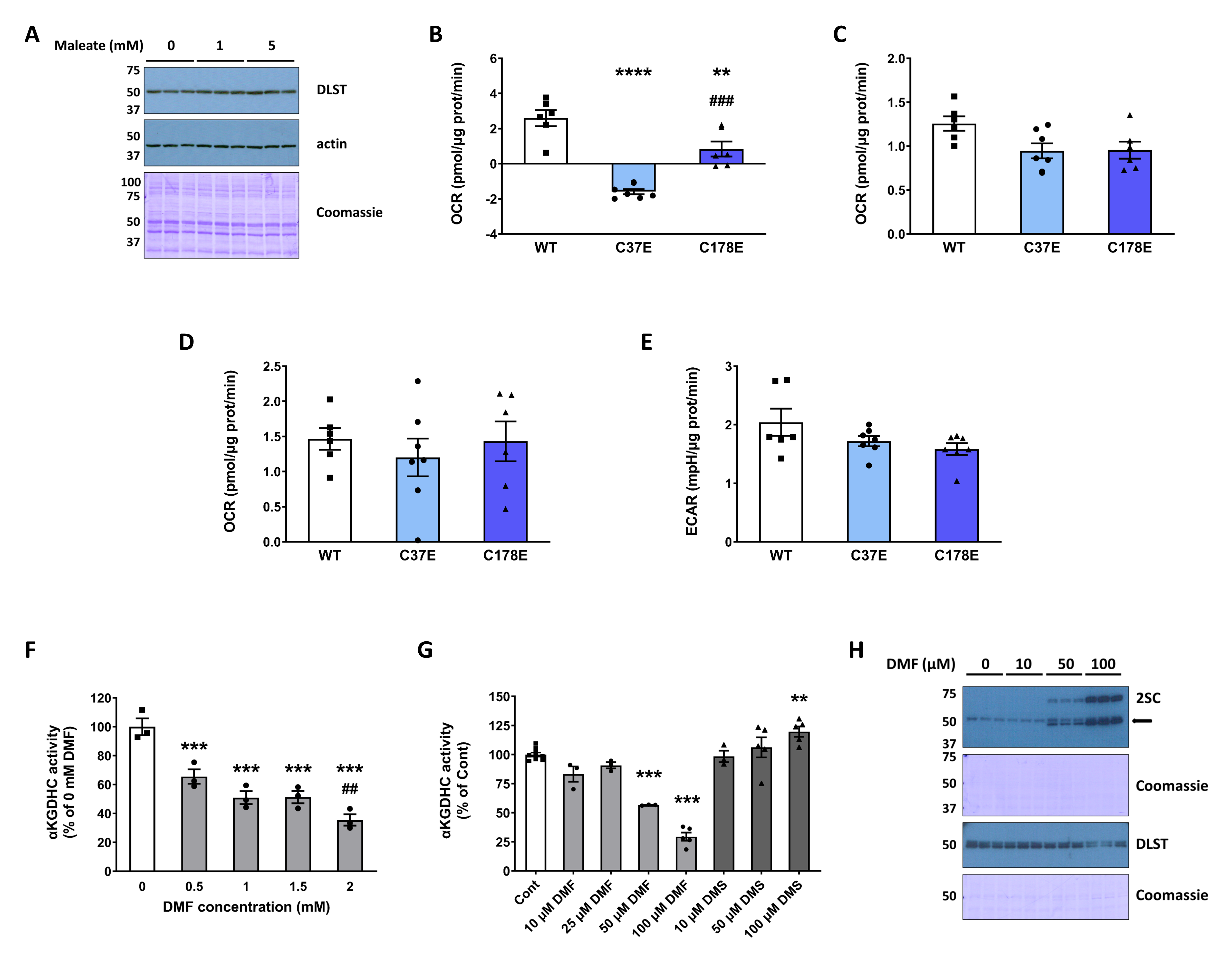

### Supplementary Figure 5

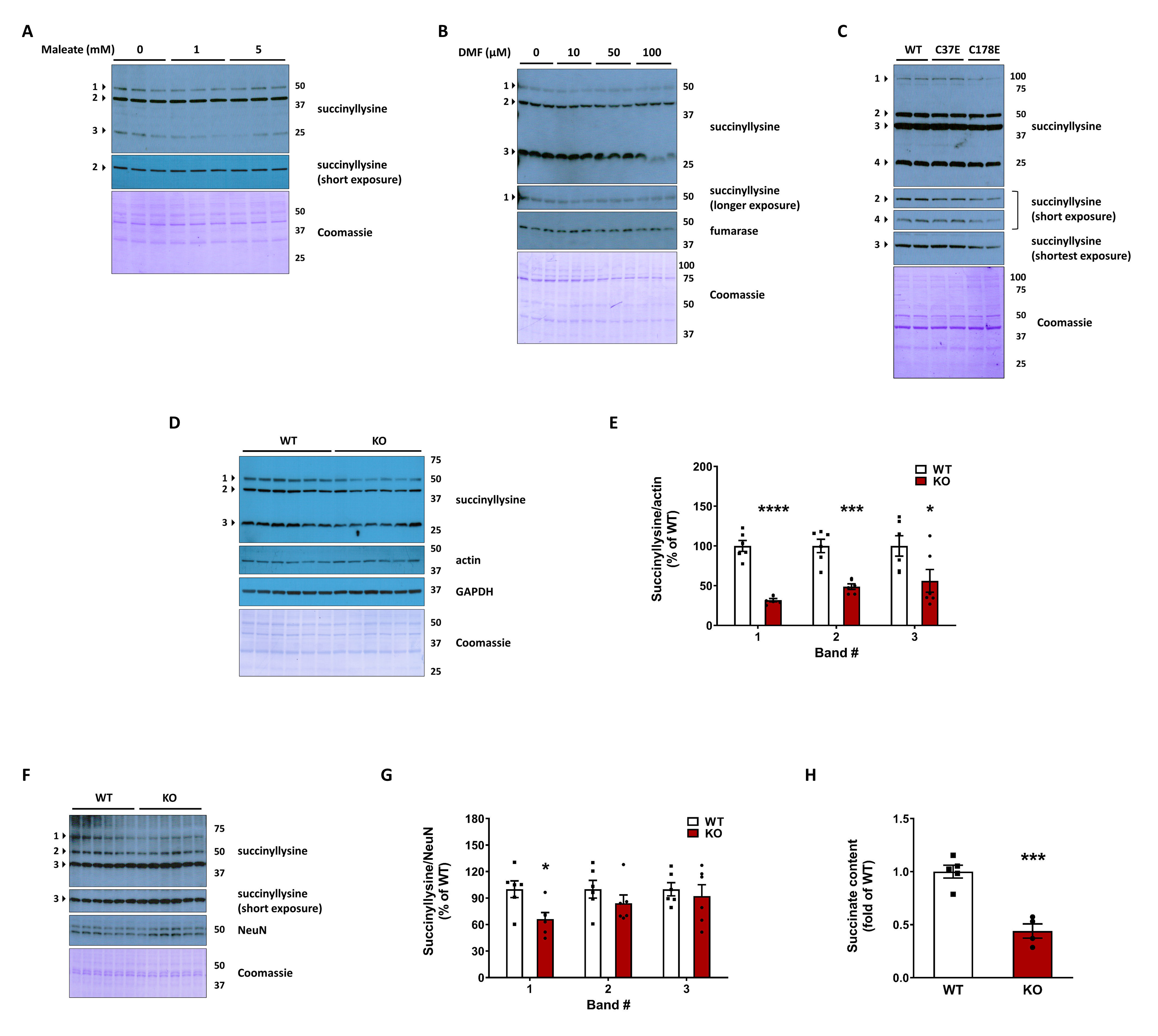
